# The use of iconPCR for 16S library preparation improves data quality and workflow

**DOI:** 10.1101/2024.12.18.629279

**Authors:** Yann Jouvenot, Caroline Obert, Brett Hale, Kyle Metcalfe, Gagneet Kaur, Andrew Boddicker, Marielle Krivet, Junhua Zhao, Tuval Ben-Yehezkel, Pranav Patel

## Abstract

Polymerase Chain Reaction (PCR) is a cornerstone of contemporary biological research, enabling the amplification of specific DNA sequences for various applications. However, suboptimal cycling conditions often undermine its efficacy, which can generate chimeric products, exacerbate PCR duplication rates, and skew species representation in metabarcoding experiments due to the preferential amplification of dominant populations.

To address these limitations, we present iconPCR—Individually Controlled PCR—a novel technology that allows each reaction in a 96-well plate to be cycled independently and programmatically. By setting a predefined fluorescence threshold, iconPCR ensures that all Next Generation Sequencing (NGS) libraries are amplified to equivalent levels, thereby eliminating the risks of over- or under-amplification through a process known as “Auto-Normalization.”

In this study, we applied iconPCR to generate V3, V4, and full-length 16S rRNA gene libraries using Avidite sequencing. The V1-V9 variable region libraries were also evaluated with HiFi sequencing, which significantly demonstrated its potential to improve microbial community analysis accuracy and reliability.

## Introduction

Polymerase Chain Reaction (PCR) is a fundamental technique in molecular biology that has revolutionized genetic research and diagnostics (Eisenstein 1990; Valones et al. 2009; Zhu et al. 2020). While advances in PCR have made it almost ubiquitous in molecular biology, certain aspects of the technology have yet to evolve since its inception. Some thermocyclers will allow users to program a gradient PCR across a plate. However, there are limitations on the temperature ranges between gradients (usually 0.5 ⁰C to 5 ⁰C), and no traditional thermocyclers allow users to vary the cycling numbers for their samples.

The iconPCR^TM^ (individually <u>con</u>trolled PCR) system is a novel real-time thermocycling instrument, developed by n6 TEC, which enables greater control of thermocycling at the single-well level. Although thermocycling protocols are programmed similarly to traditional instruments, users can also set up to 96 cycling parameters per 96-well plate. This affords users significantly greater flexibility in PCR customization. In addition to per-well temperature control, with iconPCR, the user can select a predetermined number of cycles or implement the proprietary Auto-Normalization mode (Figure 1). In the latter case, the user determines the amplified product’s fluorescence level (xBaseline), which must be achieved before final elongation and a 12°C hold. In the example shown in Figure 2, a given well’s fluorescence level must reach 2.5 times that of its baseline, which is determined as the average fluorescence of the first three cycles.

**Fig 1:**
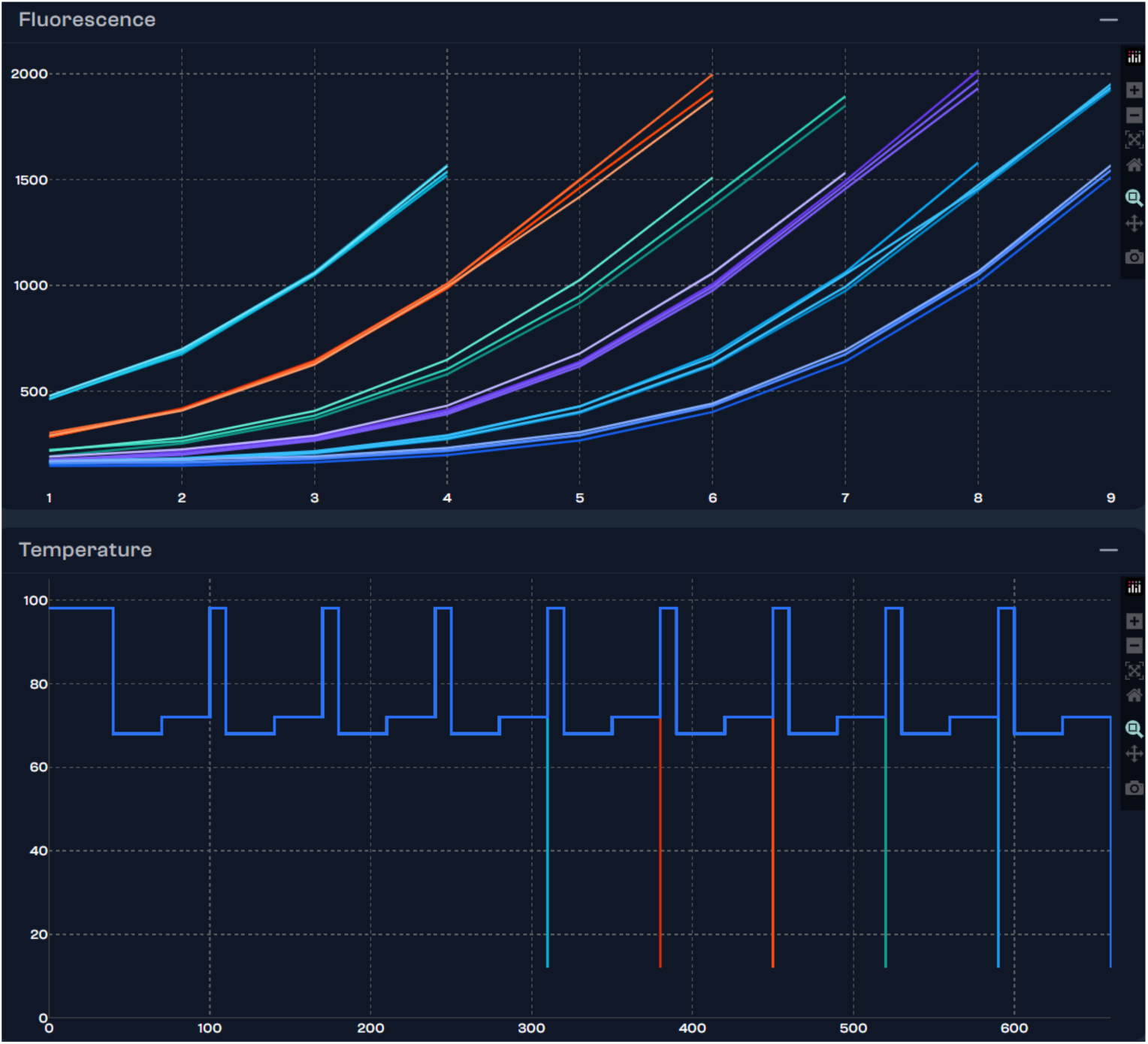
The upper panel depicts the fluorescence profiles of a serial dilution experiment (the x-axis represents cycle number, and the y-axis represents fluorescence in RFUs). The lower panel illustrates the corresponding thermal cycling profiles (the x-axis represents time in seconds, and the y-axis represents temperature in degrees C).

**Figure 2:**
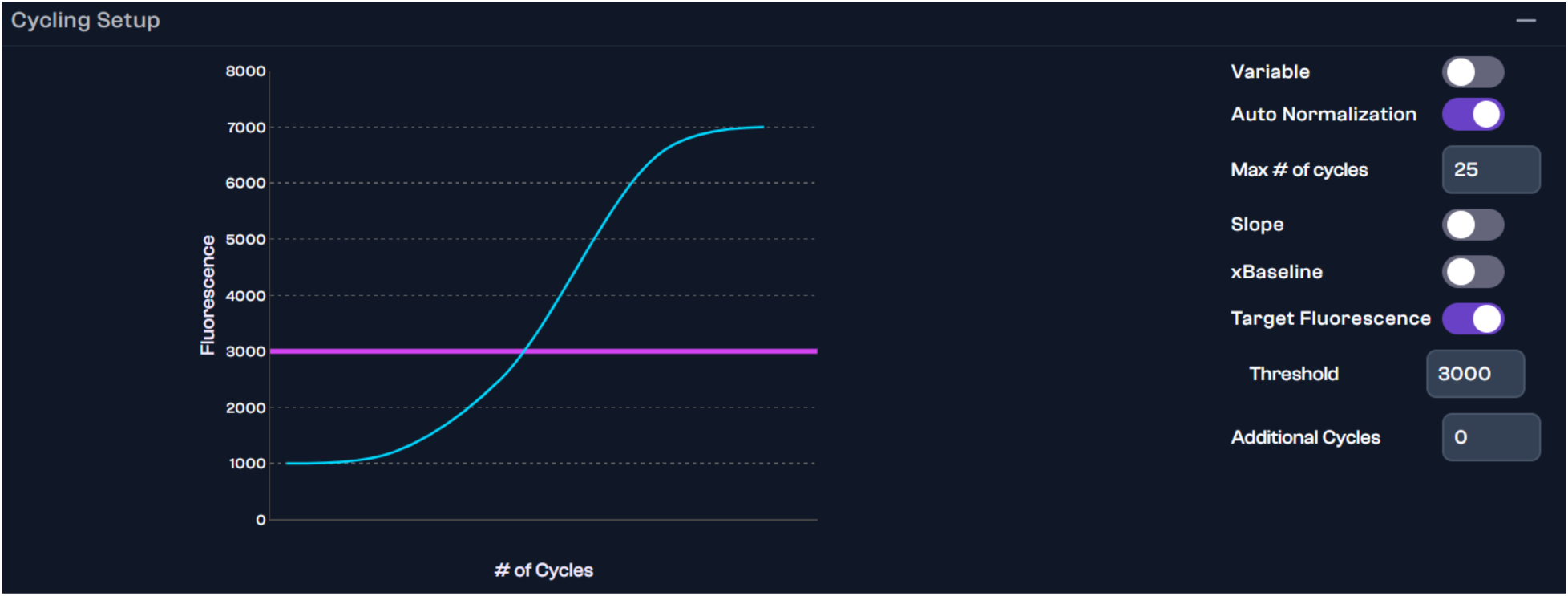
Example of Auto-Normalization settings (Target Fluorescence), In Fig 2, the blue line represents a theoretical amplification curve, while the purple line represents the threshold for Auto-Normalization (in this case, 3000 RFU).

Next Generation Sequencing (NGS) has similarly become a standard technology, especially for studying microbiome communities (Vincent et al. 2019; Wensel et al. 2022). An increasing number of applications utilize 16S rRNA gene sequencing to assess prokaryotic microbial population diversity or to detect certain pathogens (Chakravorty et al. 2017; Gilbert et al. 2014; Hajjo et al. 2022; Jin et al. 2024; Malla et al. 2019; Integrated Human Microbiome Consortium, 2014; Reese and Dunn 2018; Turnbaugh et al. 2007; Woo et al. 2008). While sequencing throughput has increased considerably over the past decade, library preparation workflows remain a significant bottleneck. Part of the issue during library preparation lies in the fact that most applications involve one or more PCR steps, and the conditions for these amplifications need to be adapted to the type of sample being studied. For example, samples with low biomass input or degraded nucleic acid quality can require additional amplification cycles to obtain the desired yield for a given assay (Witzke et al., 2020). Although the standard recommendation is to optimize the cycle number for each sample based on its quality and input amount, users will often choose the same number of cycles for all libraries, resulting in under- or over-amplification (Acinas et al. 2005; Hansen et al. 1998; Gonzalez et al. 2012; Sze and Schloss 2019).

While under-amplification will fail to produce a library that can be sequenced to a sufficient depth, over-amplification will affect the quality of the material being sequenced by creating chimeric products, exacerbating PCR duplication rates, or even altering the species representation due to the disproportionate amplification of dominant populations (Hansen et al. 1998; Gonzalez et al. 2012; Meyerhans et al. 1990; Schloss et al. 2011; Suzuki and Giovannoni 1996). The Auto-Normalization mode on the iconPCR enables users to free themselves from a predetermined PCR cycle number, thus circumventing under or over-amplification of sequencing libraries. In this study, we applied auto-normalization to generate amplicon libraries from V3, V4, and V1-V9 regions of the 16S rRNA gene and evaluated samples using Avidite and HiFi sequencing, where appropriate. In doing so, we were able to illustrate the qualitative and quantitative benefits of Auto-Normalization over Standard PCR for both short and long-read protocols in microbiome studies.

## Material and Methods

### Genomic DNA

Soil samples were collected from the rhizosphere of soybean (*Glycine max*) or field pea (*Pisum sativum*) grown under field conditions in Jonesboro, AR, USA or Fort Benton, MT, USA, respectively (Hale et al., 2024). The samples were freeze-dried immediately on solid CO_2_ and stored at −80°C until DNA isolation. The DNeasy PowerLyzer PowerSoil Kit (Cat. #12855-100, Qiagen) was used to isolate DNA from 250 mg of each sample, per manufacturer guidelines.

DNA concentrations were approximated with a Qubit fluorometer (ThermoFisher Scientific) leveraging the dsDNA high-sensitivity assay kit (Cat. #Q32851, ThermoFisher), and DNA purity was assessed from 260/230 and 260/280 nm absorbance ratios using a NanoDrop ND-1000 spectrophotometer (ThermoFisher Scientific). Negative ‘kitome’ controls were included in each isolation event.

### iconPCR Auto-Normalization

xBaseline thresholds of 2.5x were used for the Auto-Normalization conditions for all assays described below. Although the iconPCR has an upper threshold of 99 amplification cycles, all assays were normalized to a maximum of 30 cycles.

### Short Read Libraries

V3 and V4 amplicons were generated using the primers listed in Table 1, with either Standard PCR or the Auto-Normalization mode on the iconPCR, as described in Table 2 and above. Amplicons were purified using a 0.8x ratio SPRISelect beads to PCR reaction volume (B23319, Beckman Coulter) and quantified using Qubit 1x dsDNA High Sensitivity Assay (Q33231, ThermoFisher Scientific) reagents in duplicate on a Qubit 4.0 fluorometer.

**Table 1:**
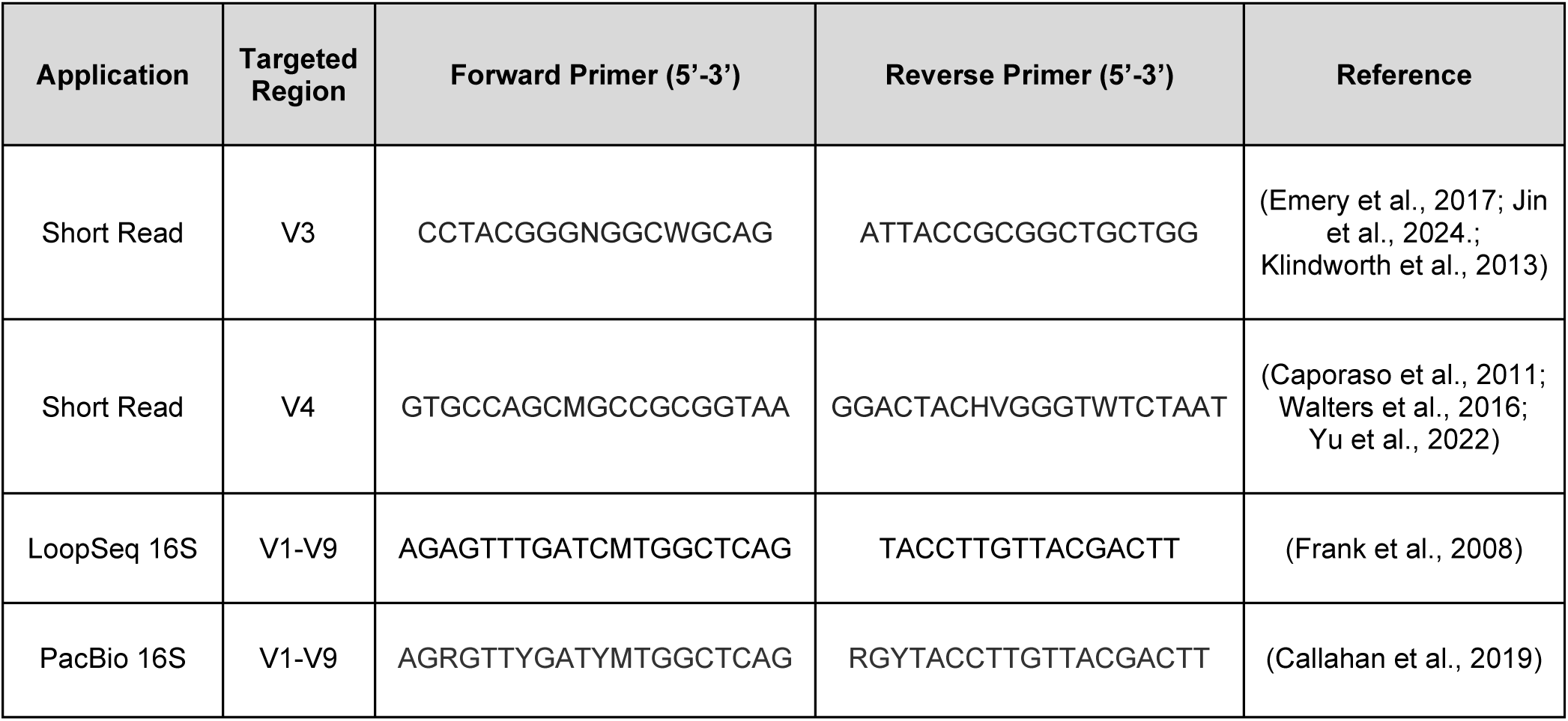
Oligonucleotide Sequences.

**Table 2:** Short Read PCR cycling conditions.

| PCR Step | Variable Region | Temperature | Time | Step |
| --- | --- | --- | --- | --- |
| Initial Denaturation |  | 98 °C | 30 s | 1 cycle |
| Denaturation |  | 98 °C | 10 s | 30 cycles or<br>Auto-Normalization |
| Annealing | V3 | 70 °C | 40 s |  |
|  | V4 | 57 °C |  |  |
| Extension |  | 72 °C | 45 s |  |
| Final Extension |  | 72 °C | 2 min | 1 cycle |
| Storage |  | 10 °C | Hold |  |

Amplicons were processed into short-read libraries using the Element Elevate Library Prep assay (Cat #: 830-00008, Element Biosciences) with Elevate Long UDI Adapter Kit Set A (Cat #: 830-00010, Element Biosciences) (Figure 3), starting with end repair and A-tailing. Insert sizes for individual samples were evaluated using the HS NGS Assay (DNF-474, Agilent) on a Fragment Analyzer 5300 system. Samples were quantitated in triplicate using Qubit 1x dsDNA High Sensitivity Assay reagents on a Synergy NEO2 plate reader and normalized to 10 ng/µL for pooling. Any sample whose concentration was too low to normalize was included by adding 10 µL of the final product. Final library pools were quantified using Qubit 1x dsDNA High Sensitivity Assay reagents in duplicate on a Qubit 4.0 fluorometer and evaluated for insert size using Agilent’s High Sensitivity DNA Kit (Cat #: 5067-4626, Agilent) for the BioAnalyzer 2100 (Agilent).

**Figure 3:**
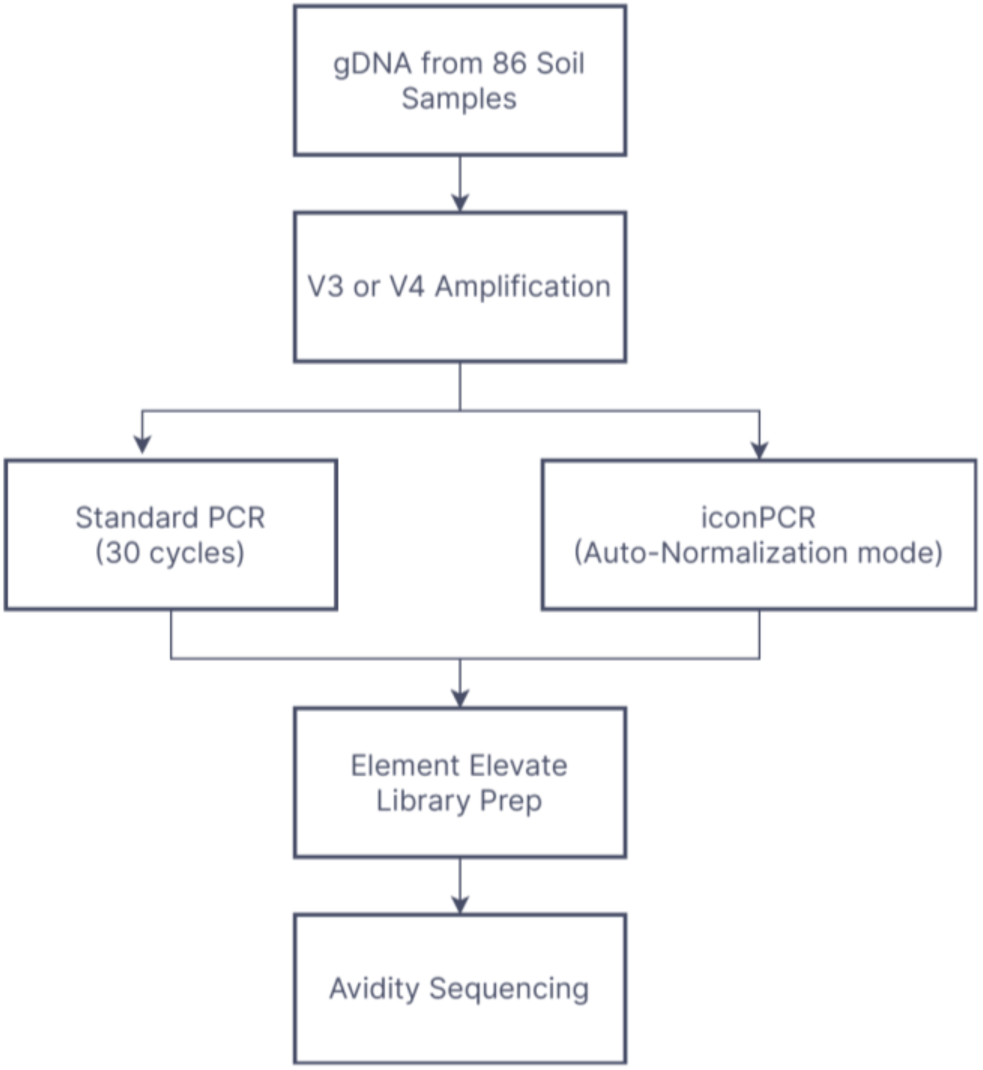
Preparation of V3 and V4 16S rRNA gene amplicons for Short Read Library Preparation

### LoopSeq Long Read 16S Libraries

Samples were prepared using LoopSeq 16S for AVITI assay (Cat. # 840-00001, Element Biosciences). ZymoBIOMICS Microbial Community DNA standard (Cat#: D6305, Zymo) was used as a control. Matched samples were processed either using Standard PCR settings, Auto-Normalization mode on the iconPCR, or with manual monitoring of amplification using the BioRad 96-well CFX Duet qPCR, as the latter platform and method has been verified for use with the LoopSeq 16S assay at Element Biosciences (Figure 4). Amplification treatments were only employed during the initial sample enrichment step of the LoopSeq 16S protocol, which used modified primers based on the sequences listed in Table 1. Samples that underwent Standard PCR cycling were amplified for 30 cycles, per the published protocol (<u>go.elementbiosciences.com/amplicon-loopseq-aviti-user-guide</u>). Monitored PCR samples underwent 22 cycles of amplification, which was determined to be sufficient as the majority of the samples had undergone 1-2 cycles of plateaued amplification. The iconPCR Auto-Normalization mode settings were as described above. All samples were subjected to melt curves after the “Enrich Samples” and “Amplify Enriched Samples” sections of the 16S protocol to evaluate for amplification artifacts. All melt curves were performed on the BioRad 96-well CFX Duet. All treatments respectively underwent 10, 24, or 14 cycles of amplification during the “Amplify Enriched Samples,” “Amplify and Pool Samples,” and “Amplify Library” sections of the protocol.

**Figure 4:**
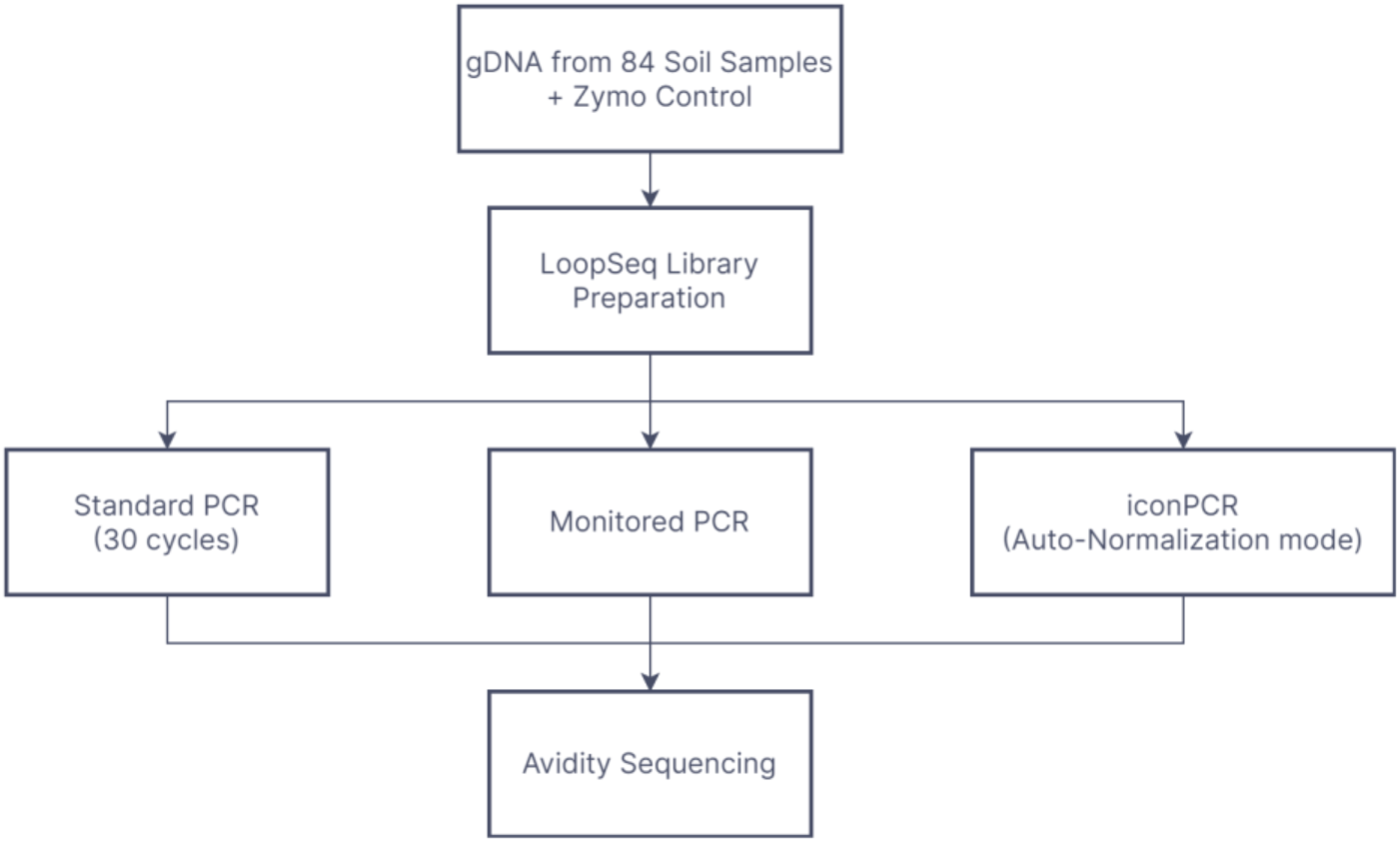
LoopSeq Long Read Library Preparation

### Avidite Sequencing

All libraries were sequenced on Element Bioscience’s AVITI platform using 2×150 Cloudbreak FreeStyle chemistry, individually addressable Lanes, and a 5% PhiX Spike-in to assist in balancing due to lower nucleotide diversity. V3 Standard PCR and iconPCR libraries were loaded at 7 pM, whereas V4 preparations were loaded at 8 pM. V1-V9 LoopSeq Standard PCR, Monitored PCR, and iconPCR libraries were loaded at 9.5 pM. Variability in loading concentrations was related to differences in final library sizes (<u>go.elementbiosciences.com/cloudbreak-sequencing-user-guide-ma-00058</u>). Preliminary demultiplexing and adapter trimming for all assays were performed using bases2fastq (https://docs.elembio.io/docs/bases2fastq/introduction/).

### PacBio Long Read 16S Libraries

A subset of sixty-five samples was evaluated for short reads, and LoopSeq synthetic long reads were prepared following the recommended PacBio protocol (https://www.pacb.com/wp-content/uploads/Procedure-checklist-Amplification-of-bacterial-full-length-16S-rRNA-gene-with-barcoded-primers.pdf) with the following modifications. ZymoBIOMICS Microbial Community DNA standard (Cat#: D6306, Zymo) was used as a control. Matched samples were processed either using Standard PCR settings or Auto-Normalization mode on the iconPCR (Figure 5).

**Figure 5:**
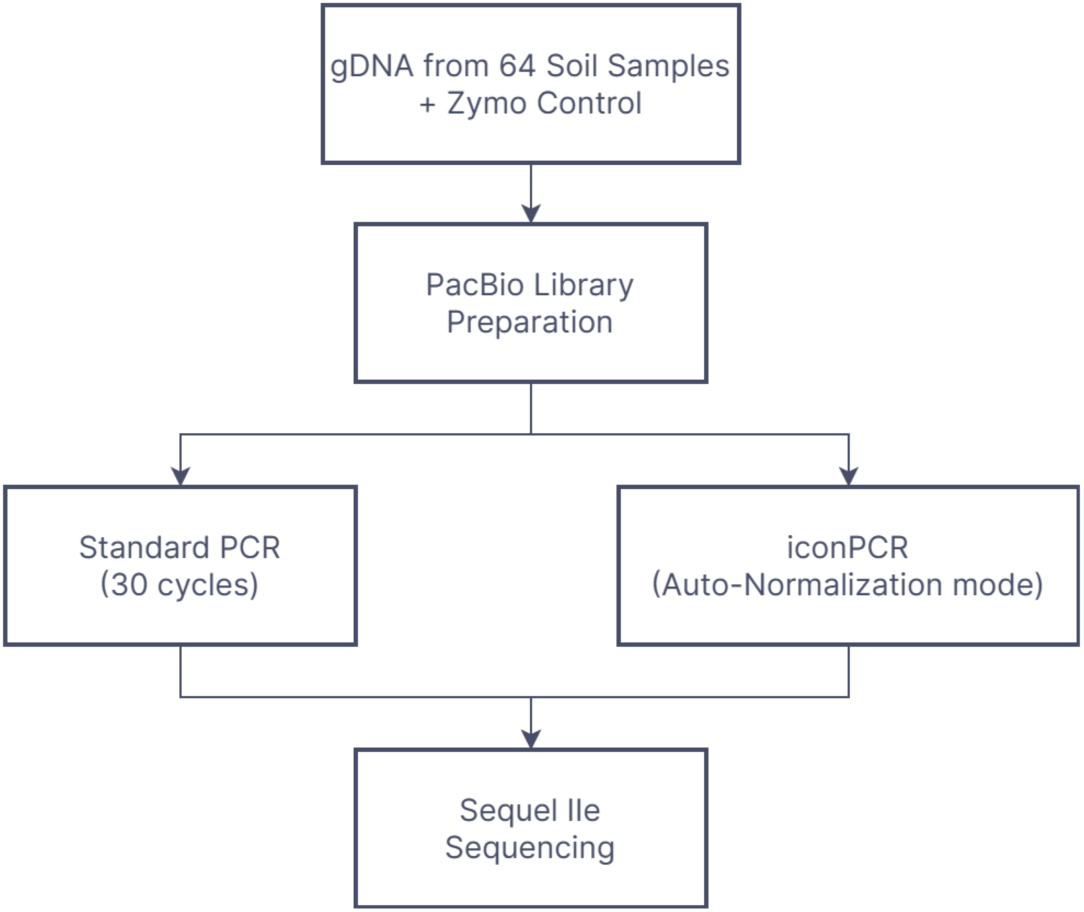
PacBio Long Read Library Preparation

Amplification for full-length 16S rRNA was achieved using modified primers listed in Table 1. Samples that underwent Standard PCR cycling were amplified for 30 cycles to reduce the number of sample dropouts. The iconPCR Auto-Normalization mode settings were as described above.

### PacBio Sequencing

All libraries were sequenced on Pacific Bioscience’s Sequel IIe system using 8M SMRT cells.

## Data Analysis

### Qualitative Analysis - Short Amplicons and LoopSeq

Final V3 and V4 libraries averaging ∼314 bps and ∼422 bps were expected based on initial amplicon sizes of 175 bps and 283 bps, respectively (Vargas-Albores et al., 2017). LoopSeq Library prep melt curves were assessed in the CFX Maestro Software 2.3 (version 5.3.022.1030). Library insert sizes were evaluated either with ProSize Data Analysis Software (5.0.1.3) or 2100 Expert Software (Version B.02.11SI824 [SR1]).

### Synthetic Long Read Assembly

Consensus sequences from the LoopSeq application were assembled via the LoopSeq Analysis pipeline (Liu et al., 2021), a version of which is freely available on GitHub (https://github.com/Elembio/loop-core). In brief, after samples are demultiplexed by the pool index on the sequencer, they undergo adapter sequence trimming via Trimmomatic (Bolger, 2014). They are then subsequently demultiplexed by plate positional index and UMI. The binned UMIs are de novo assembled using SPAdes (Bankevich, 2012). Individual short reads, which may have sequencing or chimera-related errors, are evaluated during the SPAdes assembly of the synthetic long read, increasing the final sequence’s overall accuracy.

### Chimera Analysis - Short Read and LoopSeq Workflows

Chimera rate analysis was performed using DADA2 v1.30.0 in R v4.3.0 (Callahan et al., 2016). In brief, LoopSeq synthetic long reads, as well as V3 and V4 short reads, were deduplicated, and ASV (amplicon sequence variant) composition was inferred using DADA2 with default parameters. Chimeras were detected using the DADA2 “removeChimeraDenovo” function (Prodan et al., 2020). Values of zero indicate that the software did not detect chimeras in the sample, while the rate calculated indicates the fraction of reads that were determined to be chimeras.

The normality of the chimera rate distribution was tested for each amplicon-cycling method combination using the Shapiro-Wilk test (Shapiro and Wilk, 1965) implemented with the ‘shapiro.test’ function of the *stats* package (v4.4.0) in RStudio (v4.2.2) (Racine, 2012). The Shapiro-Wilk test suggested non-normality in all instances (p < 0.05); thus, the Wilcoxon signed-rank test was leveraged to discern differences among cycling method population mean ranks with the *stats* function ‘wilcox.test’ (paired = TRUE). Chimera rates and statistical outputs were visualized with *ggplot2* (v3.4.2) (Wickham, 2011).

### Chimera Analysis - PacBio Workflow

The DADA2 pipeline inferred unique ASVs from raw reads (Callahan et al., 2016) and removed potential sequencing errors and chimeric sequences.

### Short Read Taxonomic Analyses

Taxonomy was assigned to non-chimeric ASVs using the DADA2 “AssignTaxonomy” function, which uses the SILVA 138.1 database (Pruesse et al., 2007) with default parameters (https://zenodo.org/records/4587955).

### Long Read Taxonomic Analyses - LoopSeq

Taxonomy was assigned via QIIME2 (v2024.2.0) (Caporaso et al., 2010) with the SILVA 138.1 database clustered at 99% identity. Shannon Index (Shannon, 1948) alpha-diversity analyses were performed with QIIME2 (v2024.2.0). A diversity value equal to one indicates that all species within a sample had the same frequency. A diversity value equal to zero indicates that only one species was detected.

### Long Read Taxonomic Analyses - PacBio

Taxonomy assignment was performed using Uclust from QIIME v.1.9.1 with the proprietary Zymo Research 16S Database. Shannon Index alpha-diversity analyses were performed with QIIME v.1.9.1. Values could not be calculated for any sample with less than 4000 reads after DADA2 read processing.

### Statistical Analysis

Matched chimeric rates and Shannon indexes were evaluated for normality using the Anderson-Darling test (Anderson & Darling, 1954), equality using non-parametric Friedman (Friedman, 1937, 1939) or Wilcoxon (Wilcoxon, 1945) tests, as appropriate, and equivalency using the Bland-Altman Test (Bland and Altman, 1999; Bland and Altman 2003) and Student’s T-test (Student, 1908). The Zymo mock community comparisons were evaluated using Spearman Rank Correlation (Spearman, 1904; Shetty et al., 2023). All analyses were calculated using the Analyse-It 6.15.4 statistical package.

## Results and Discussion

### Cycling results

Although the advantages of reduced amplification cycles in the generation of 16S amplicons have been widely touted (Ahn et al., 2012; Bonnet et al., 2002; Pollack et al., 2018; Qiu et al., 2001; Suzuki and Giovannoni, 1996; Wu et al, 2010), the benefits of including low biomass samples (through increased cycling) to better understand microbial communities has recently garnered traction within the microbiology community (Karstens et al, 2018; Witzke et al, 2020).

The iconPCR system in Auto-Normalization mode used 19 to 23 amplification cycles for the V3 libraries, 11 to 18 cycles for the V4 libraries, 7 to 30 cycles for LoopSeq libraries, and 17-21 cycles for PacBio samples. The variation in cycle numbers reflects the differences in sample input amounts, reaction efficiency, and process variations (e.g., variability in pipetting over the course of preparation or differences between library prep protocols). Increased amplification cycles for low biomass samples were expected with long-read library applications. Synthetic long-read applications require numerous copies of the initial long molecules to provide the error-corrected consensus sequences. Although sixty-four samples were initially processed using the PacBio assay, matched data was only available for thirty-three samples. The majority of the samples that could not be processed completely due to low read generation (<10k per sample) after processing through the proprietary Zymo pipeline were Standard PCR preparations on a single plate, suggesting a failure during library preparation. As such, the samples were excluded from subsequent downstream analyses. Auto-Normalized cycle numbers per well for V3 and V4 are available in Supplemental Table 1 and for both full-length assays in Supplemental Table 2.

### Library Quality

Evaluation of NGS libraries, either over the course of the preparation (as in the case of LoopSeq libraries) or at the end of preparation, can reveal the formation of undesirable artifacts. Smaller products (<150 bps) are often adapter or primer dimers, while larger products are generally chimeras caused by incomplete PCR products serving as primers to related sequences (Schloss et al., 2011). The presence of these molecules in a final library pool can skew the amplification and resulting sequencing of final library preparations.

The figures below compare Bioanalyzer profiles between matching sample pools prepared by Standard PCR protocol with fixed cycles (FC) or using Auto-Normalization (AN). The red electropherogram in Figure 6 represents the final library pool from V3 amplicons that underwent Standard PCR. In that trace, there is a dominant peak around 1200 bp, while the expected product V3 amplicon (300-350 bp) constitutes a much smaller fraction of the pool (9% of the total pool). The blue electropherogram, representing the V3 iconPCR amplicon pool, only shows a double peak (∼78% of the total pool) around the expected sizes. Similarly, the top panel in Figure 7 depicts the overlay of the final V4 amplicon pools. The V4 Standard PCR pool (in red) has various peaks greater than 500 bp, with only 19% of the total pool in the expected region.

**Figure 6:**
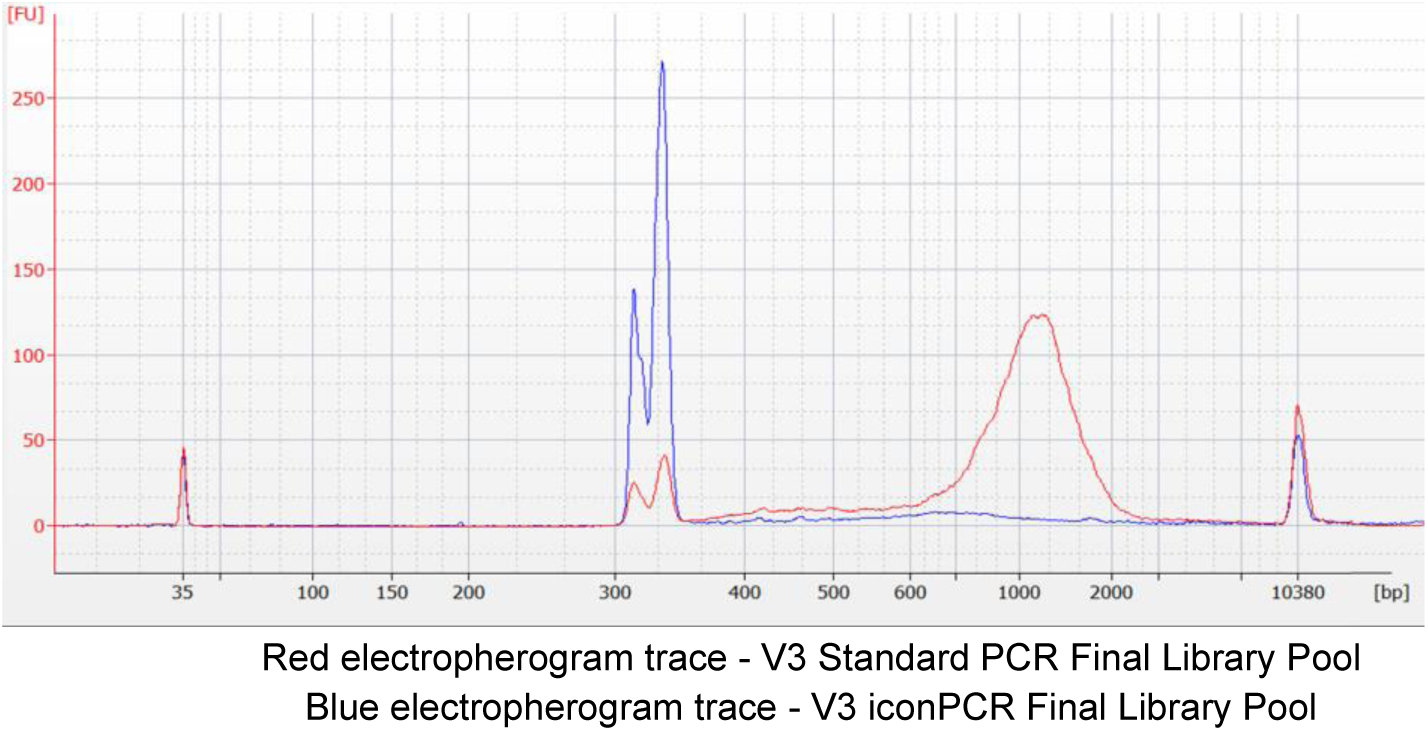
Comparison of Bioanalyzer Profiles for Final V3 Library Pools.

**Figure 7:**
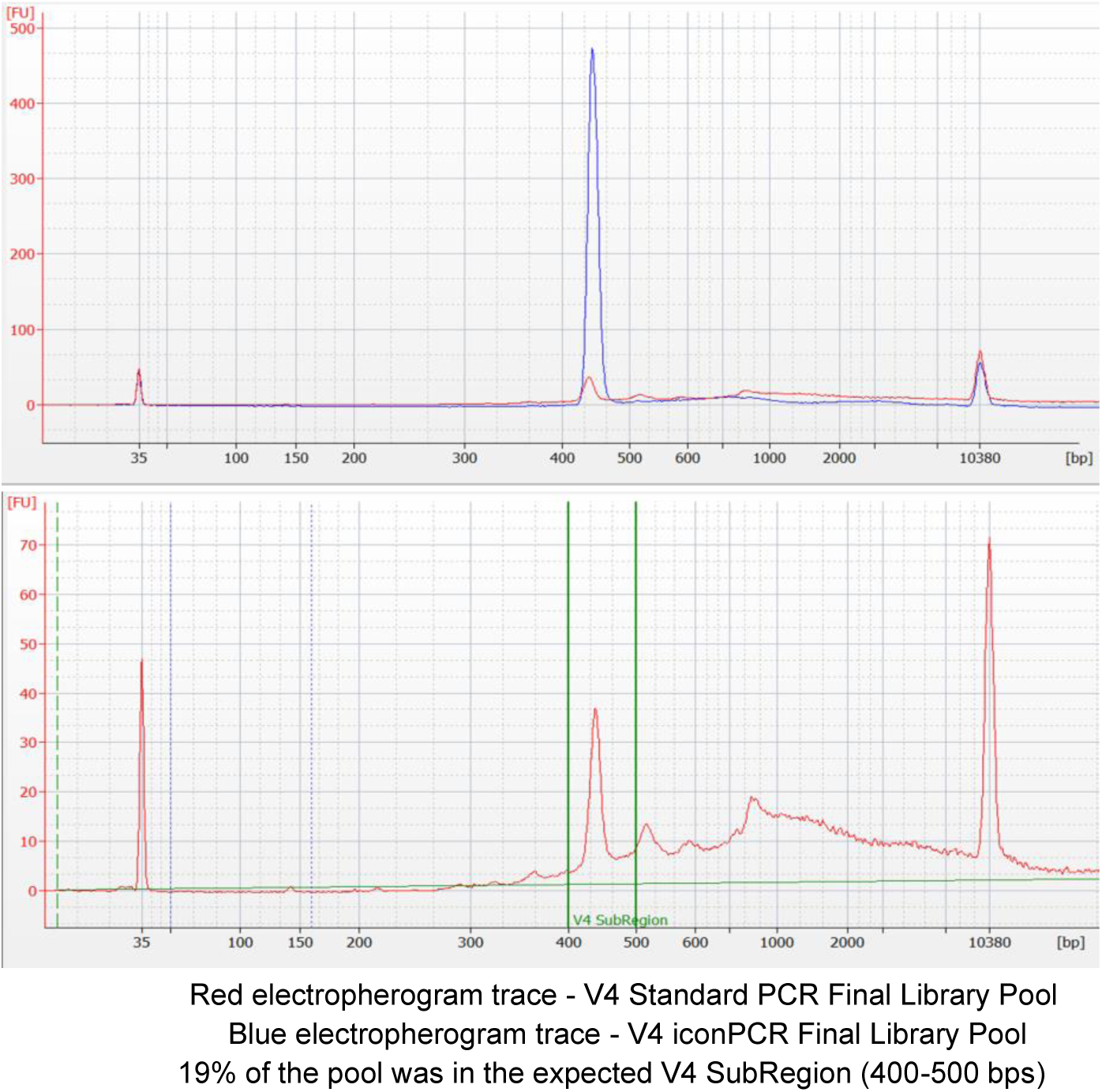
Comparison of Bioanalyzer Profiles for Final V4 Library Pools.

This trace is shown in greater resolution in the lower panel, without the overlay. By comparison, the V4 iconPCR pool (in blue) only displays one peak around the expected size (∼440 bp) for the matched pool (74% of the total pool) generated using Auto-Normalization. Similar findings were observed in the LoopSeq preparations after enrichment of the full-length 16S product, where products >1500 bps were significantly more prominent in Monitored PCR and Standard PCR preparations compared to Auto-Normalized samples (data not shown).

### Chimeric Analysis

In 16S analysis, chimeras refer to artificial DNA sequences created during PCR amplification, where parts of two different bacterial DNA sequences are joined together, resulting in a sequence that appears to be from a novel organism. If the sequences are not identified and removed bioinformatically after sequencing, one risks potentially distorting the results of microbiome analysis by inflating perceived diversity and leading to incorrect taxonomic assignments (Pollack et al., 2018; Witzke et al., 2020).

A bioinformatic analysis of the sequencing data uncovered a statistically significant difference (p<0.0001) in chimera rates between Standard PCR and iconPCR libraries for V3 and V4 libraries (Figure 8) when evaluated using the Wilcoxon Test at 5% significance (Supplemental Tables 3 and 4). Of the 83 samples for which there was matched data, 75 (90.4%) of the V3 amplicon samples had lower average chimera rates (and inversely higher percentages of chimera free-reads) using the iconPCR Auto-Normalized mode (0.031515 ± 0.001275 (SE)) instead of Standard PCR (0.056335 ± 0.000349). No specific factors were identified (e.g., sample origin, sample input concentration, or cycling number) among the eight samples with lower chimera rates/higher chimera-free reads with Standard PCR, which account for the observed deviation. Amongst the 86 samples with matching data, 100% of the V4 amplicon samples had lower chimera rates and a higher percentage of chimera free-reads using Auto-Normalized mode (0.092102 ± 0.011606 (SE)) when compared to Standard PCR (0.396882 ± 0.009988 (SE)). Individual sample chimera rates for avidite-prepared libraries are provided in Supplemental Tables 3-4.

**Figure 8.**
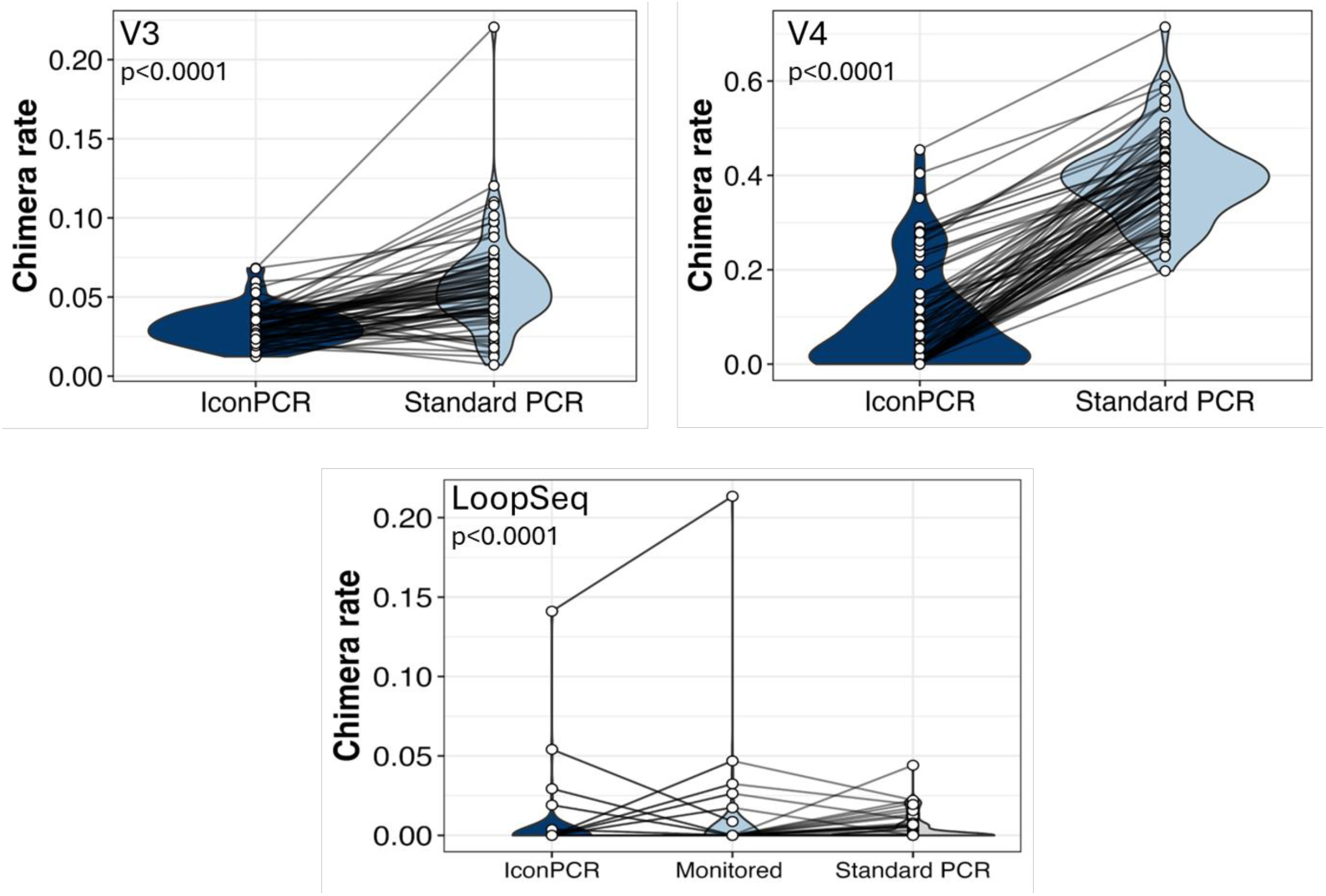
Chimeric Rate Comparisons

The Loop Core pipeline corrects for chimeric short reads during the error-correction de novo assembly of the synthetic long-reads. While this does not eliminate the presence of all chimeras, it significantly reduces the number of final reads later identified as chimeric. The Friedman test analysis of chimeric rate equalities among treated samples processed through the LoopSeq assay (Figure 8) indicated a significant difference (p<0.0001 at 5% significance) between treatments. The average chimeric rates for iconPCR, Monitored PCR, and Standard PCR were 0.003053 ± 0.001885 (SE), 0.004061 ± 0.002622 (SE), and 0.004864 ± 0.001046 (SE), respectively. However, the differences are attributable to the Standard PCR treatment, as iconPCR and Monitored PCR chimeric rates were statistically equal (p=0.4961, using a Wilcoxon test at 1% significance).

As different pipelines were employed to evaluate chimerism across methods, a comparison of the calculated chimera-free reads was employed to evaluate the impact of iconPCR Auto-Normalization (Figure 9). In each application, the implementation of the Auto-Normalization mode resulted in a significantly higher (p<0.0001) percentage of chimera-free reads when compared to Standard PCR. The limited influence of chimera generation across LoopSeq applications is related to the per-base error-correction analyses employed in the Loop Core pipeline (Callahan et al., 2021). Although the V3 libraries demonstrated the generation of disparately larger chimera products (Figure 6), it is likely that relatively fewer of those sequences underwent polony generation and rolling circle amplification on the AVITI platform (Arslan et al., 2024), resulting in a less pronounced difference in chimera-free reads between treatments. However, the effect of the Auto-Normalization mode is evident by the significant differences in the PacBio preparations and V4 short read amplicon (46% and 30%, respectively). Given that the V4 amplifications generated relatively more similar-sized chimeras (Figure 7), the sequencing of more chimeras was expected. Individual sample chimera-free read values are summarized in Supplemental Tables 3-6.

**Figure 9.**
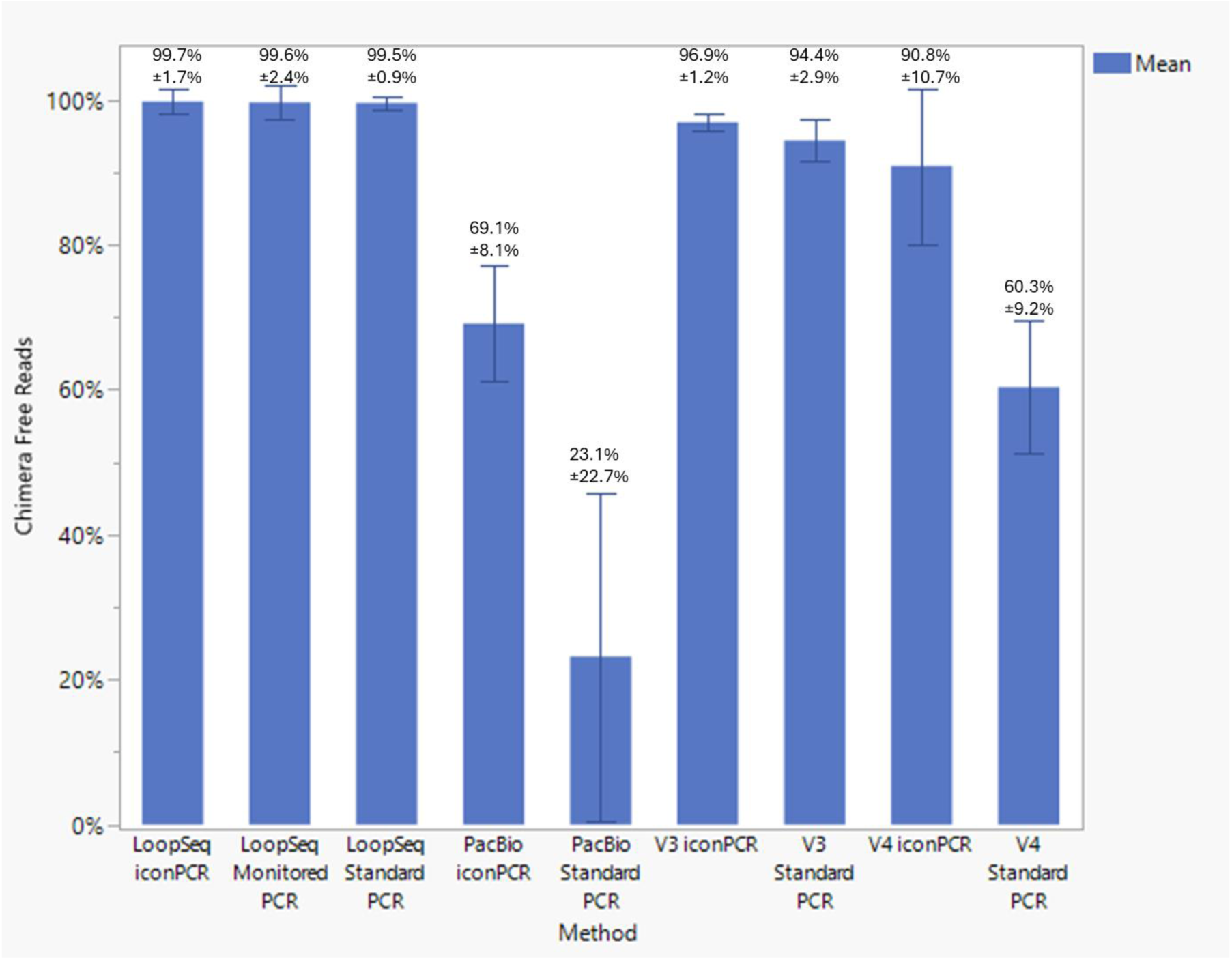
Chimera-Free Reads

### Detected Species Diversity

To avoid concerns regarding variability in species detection when using sub-regions (e.g., V3 or V4) of the 16S rRNA gene (Abellan-Schneyder et al., 2021; Chung et al., 2020; Jeong et al., 2021; Yu et al., 2022), species diversity was evaluated for full-length V1-V9 application using orthogonal assays. LoopSeq 16S preparations were evaluated for the number of full-length molecules between 1276-1726 bps in length, as smaller and larger fragments generally failed to be assigned an ASV value through the LoopSeq 16S pipeline. iconPCR Auto-Normalized molecules averaged 1474 ± 111 bps, of which 98.9% fell into the size range, whereas Monitored PCR molecules averaged 1471 ± 104 bps, with 98.4% of the molecules within the range (Table 3). By comparison, 30 cycles of Standard PCR amplification resulted in molecules averaging 1052 ± 552 bps in length, with only 61.7% of the molecules ascribed to the size range. Although iconPCR PacBio samples had a relatively higher percentage of ASVs (Table 3) and detected species (Figure 10) for matched samples over Standard PCR, the marked difference in full-length product lengths was not observed. PacBio samples that were processed using iconPCR averaged 1490 ± 26 bps, while standard PCR samples averaged 1495 ± 26 bps. The differences in full-length sizes are likely attributable to the inherent differences in library preparation for synthetic long reads versus native long reads.

**Figure 10.**
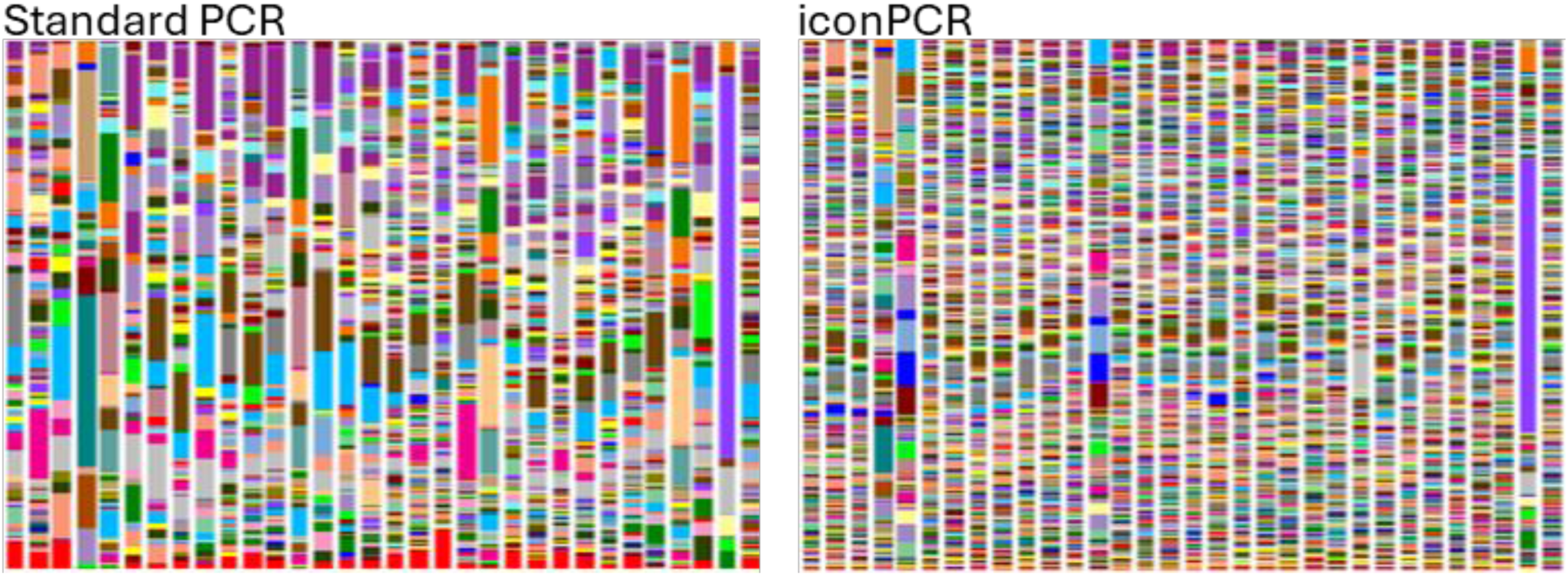
PacBio Species Diversity

**Table 3.** Long Read Molecules and Detected ASVs by Treatment.

| Treatment | Samples Evaluated | Total Full-Length Molecules (bp) | Avg. Molecule Length (bp) | Full-Length Molecules Between 1276-1726 bps (%) | Unique ASVs (%) | ASVs without Singletons (%) | ASVs without Doubletons (%) |
| --- | --- | --- | --- | --- | --- | --- | --- |
| LoopSeq iconPCR | 85 | 253,102 | 1474 ± 111 | 250,223 (98.9%) | 234,023 (92.5%) | 7,349 (2.9%) | 3,220 (1.3%) |
| LoopSeq Monitored PCR | 85 | 421,512 | 1471 ± 104 | 414,866 (98.4%) | 412,518 (97.9%) | 3,386 (0.80%) | 1,409 (0.33%) |
| LoopSeq Standard PCR | 85 | 150,059 | 1052 ± 552 | 92,629 (61.7%) | 87,653 (58.4%) | 1,472 (0.98%) | 434 (0.29%) |
| PacBio iconPCR | 33 | 1,681,760 | 1490 ± 26 | 1,679,763 (99.8%) | 991,488 (59.0%) | N/A | N/A |
| PacBio Standard PCR | 33 | 780,585 | 1495 ± 26 | 779,049 (99.8%) | 188,342 (24.1%) | N/A | N/A |

The percentages of ASVs listed represent the fraction of the total full-length molecules that were classified as ASVs. ASV values differ from Total Full-Length Molecule values because they represent the unique sequences observed in the pool rather than the total number of sequences.

When ASV richness was assessed across cycling treatments (Table 3), Monitored PCR had a marginally higher percentage (97.9%) of ASVs than iconPCR (92.9%). However, when singleton and doubleton sequences were removed, it comparatively had the lowest number of classified ASVs (0.8% and 0.33% vs 2.9% and 1.3%, respectively). This marked difference, especially with regard to LoopSeq iconPCR, is likely related to the over-cycling of samples while visually determining when the majority of the samples on the plate had achieved a desired cycling amplification value.

The Zymo Microbial Community Standard was included as a control in each method tested. Figure 11 illustrates the community composition detected for each different method. Table 4 evaluates the distribution of the eight bacterial species included in the sample using the Spearman rank sum correlation. The results indicate that while there was no notable statistical difference between the distributions for Monitored LoopSeq and iconPCR (r_s_ = 0.952), the former method was slightly more correlated to the expected distribution (r_s_=0.857 vs. r_s_= 0.810, respectively). The lowest correlations were associated with Standard PCR methods, regardless of the assay, with Standard LoopSeq having a marginally improved correlation over Standard PacBio (r_s_= 0.491 vs. r_s_= 0.431, respectively). Overall, this comparison suggests that reducing overamplification in conjunction with the LoopSeq assay may provide a more accurate representation of species abundances.

**Figure 11.**
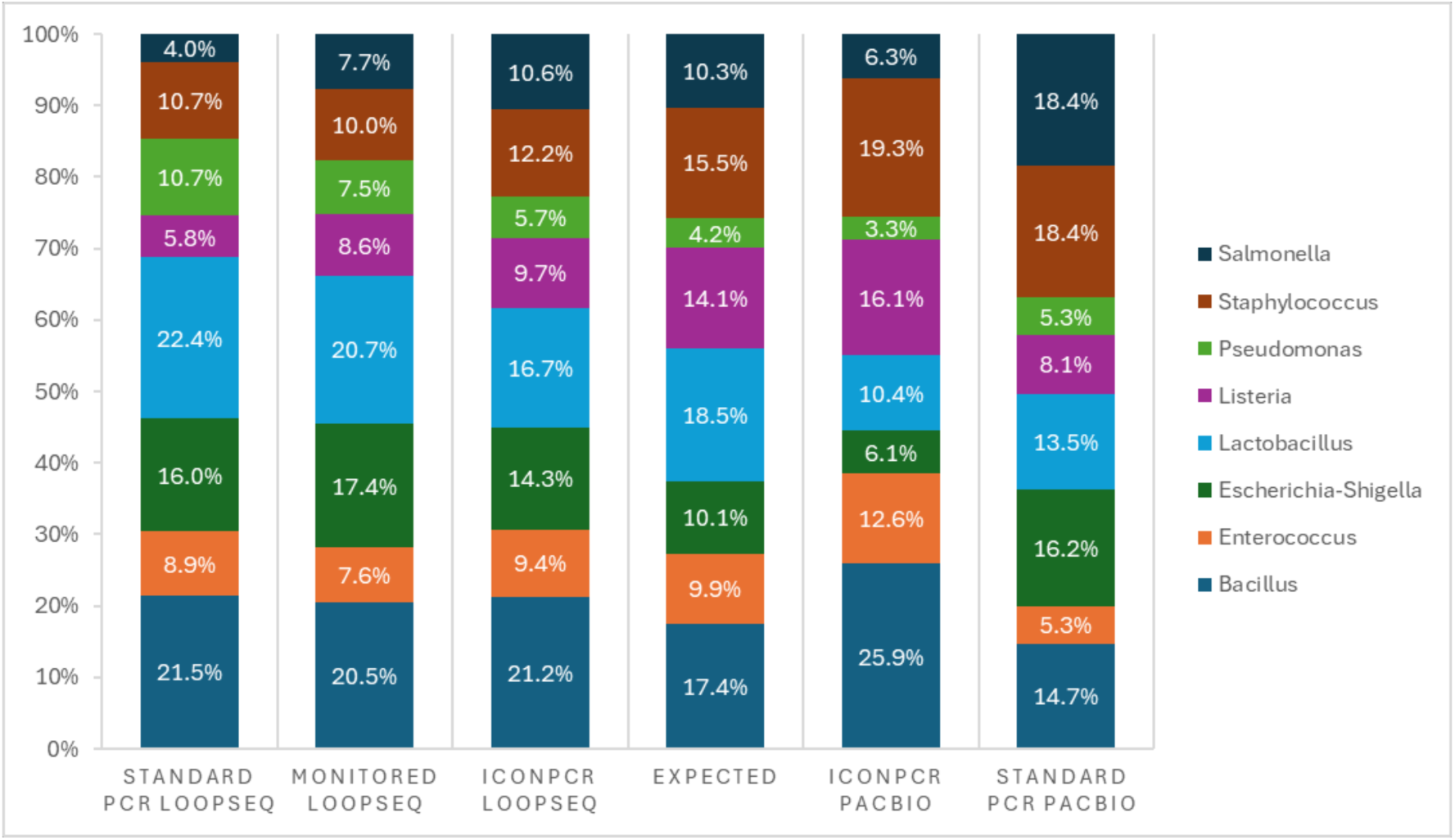
Zymo Microbial Community Standard

**Table 4.**
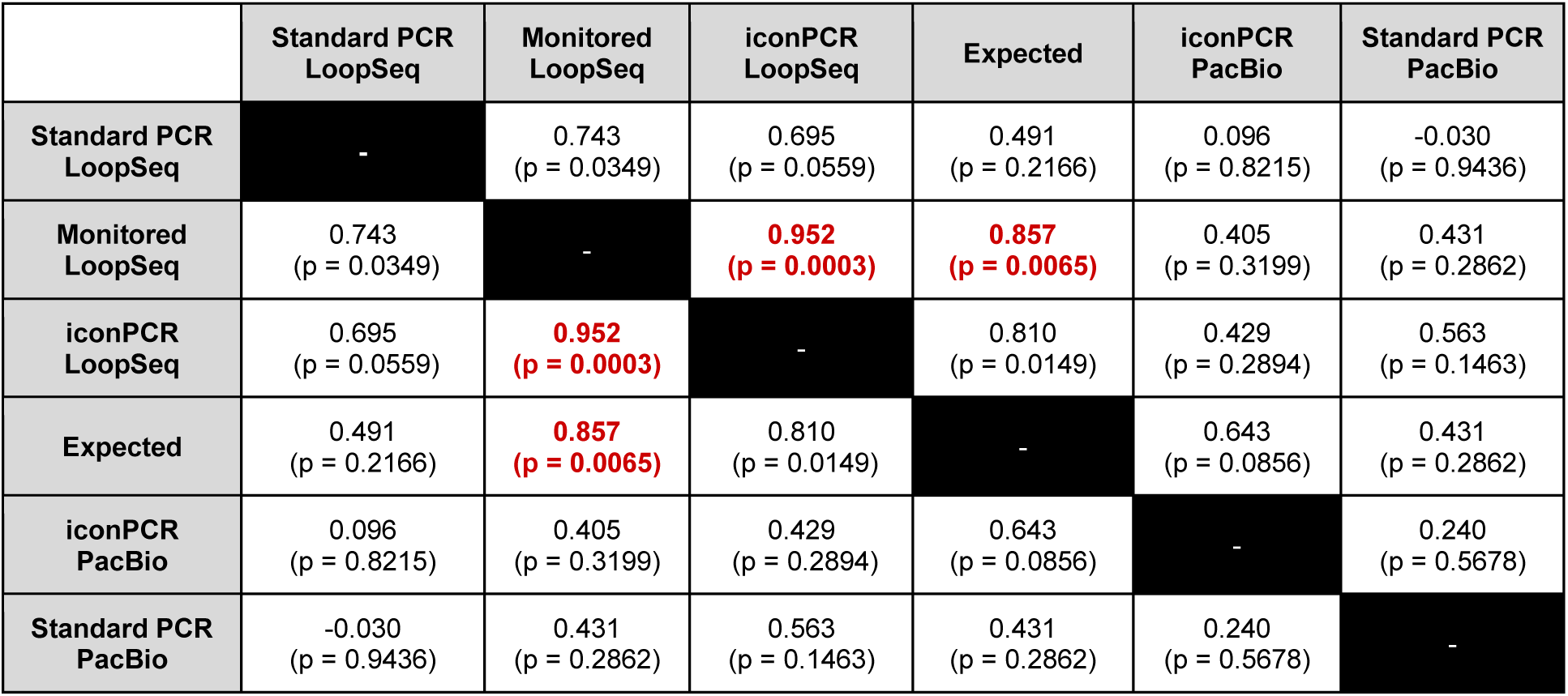
Spearman Rank Sum Correlation of Zymo Mock Community. Bolded red values indicate that no statistical difference was observed between conditions.

|  | Standard PCR LoopSeq | Monitored LoopSeq | iconPCR LoopSeq | Expected | iconPCR PacBio | Standard PCR PacBio |
| --- | --- | --- | --- | --- | --- | --- |
| Standard PCR LoopSeq | - | 0.743<br>(p = 0.0349) | 0.695<br>(p = 0.0559) | 0.491<br>(p = 0.2166) | 0.096<br>(p = 0.8215) | -0.030<br>(p = 0.9436) |
| Monitored LoopSeq | 0.743<br>(p = 0.0349) | - | **0.952**<br>(p = 0.0003) | **0.857**<br>(p = 0.0065) | 0.405<br>(p = 0.3199) | 0.431<br>(p = 0.2862) |
| iconPCR LoopSeq | 0.695<br>(p = 0.0559) | **0.952**<br>(p = 0.0003) | - | 0.810<br>(p = 0.0149) | 0.429<br>(p = 0.2894) | 0.563<br>(p = 0.1463) |
| Expected | 0.491<br>(p = 0.2166) | **0.857**<br>(p = 0.0065) | 0.810<br>(p = 0.0149) | - | 0.643<br>(p = 0.0856) | 0.431<br>(p = 0.2862) |
| iconPCR PacBio | 0.096<br>(p = 0.8215) | 0.405<br>(p = 0.3199) | 0.429<br>(p = 0.2894) | 0.643<br>(p = 0.0856) | - | 0.240<br>(p = 0.5678) |
| Standard PCR PacBio | -0.030<br>(p = 0.9436) | 0.431<br>(p = 0.2862) | 0.563<br>(p = 0.1463) | 0.431<br>(p = 0.2862) | 0.240<br>(p = 0.5678) | - |

Species-level taxonomic identification of soil samples remains limited due to long-standing issues pertaining to bacterial culturing that are only now being addressed through technological improvements in sequencing (Zhang and Xu, 2008). However, while sequencing advancements have been made, the accurate classification of presumably rare strains (represented by singletons or doubletons in the data) is elusive. The utility of the inclusion of singletons and doubletons in the assessment of species diversity is often debated. While many view their presence to be the result of sequencing error whose inclusion overinflates estimates of species richness (Callahan et al., 2021; Chiu and Chao, 2015), recent studies by Wells et al. (2019) have also shown that in previously under-sampled environments it is possible to resample the communities and discover additional samples which support the validity of previously ascribed rare sequences. These findings have led to methods for correcting the biased data to obtain a more accurate representation of the microbial diversities in the presence of rare sequences (Gatti et al., 2020; Yen and Chiu, 2020).

Although we refrained from employing either of the accuracy correction methods noted above, we opted to assess the impact of cycling treatments on detected species diversity by comparing the number of detected ASVs and the corresponding Shannon Diversity Indexes across all matched LoopSeq prepared samples when all error-corrected sequences were included (Figure 12) and when singletons (Figure 13) or doubletons (Figure 14) were removed (Supplemental Tables 7 and 8).

**Figure 12.**
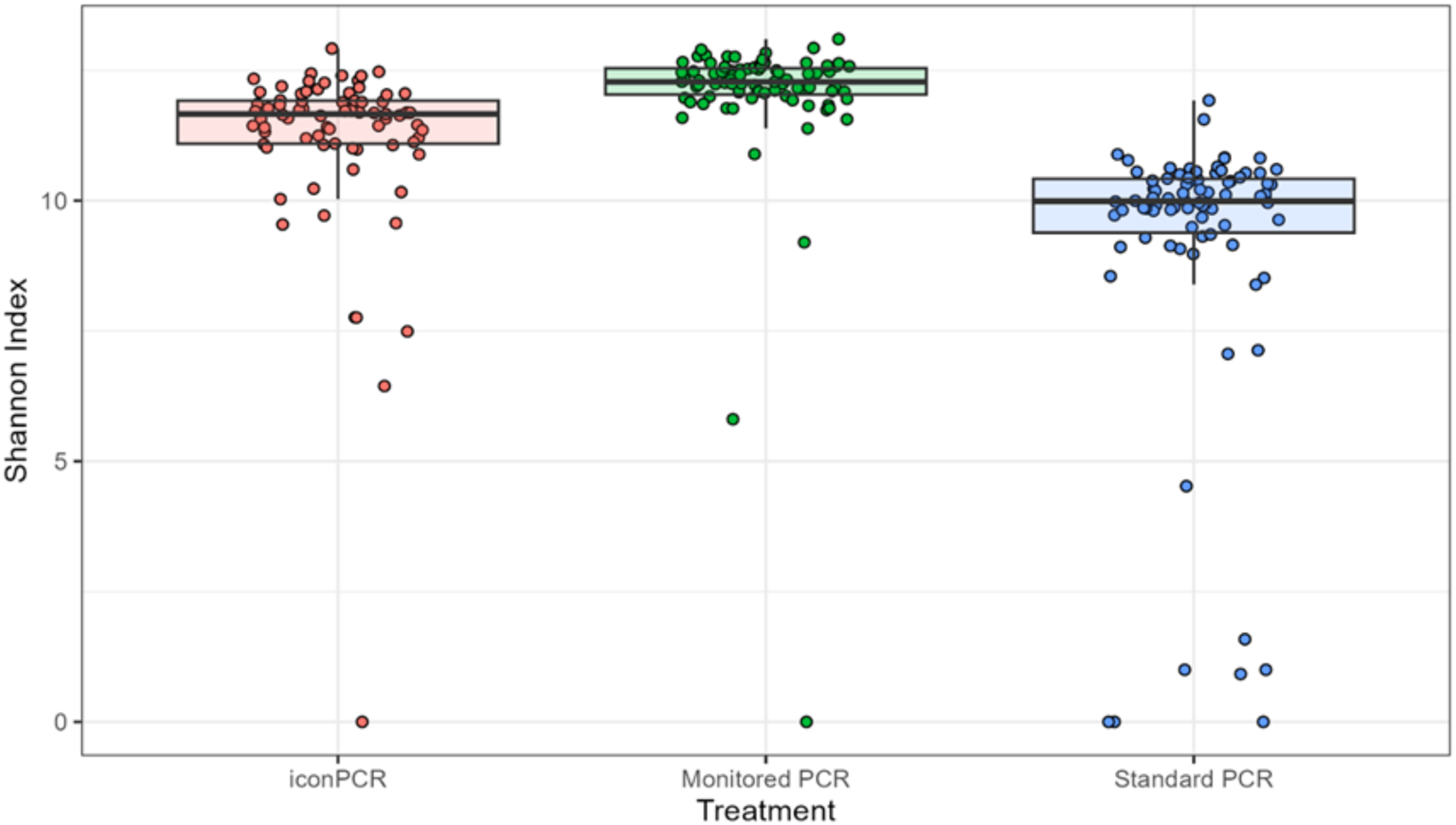
Detected Diversity by Treatment - LoopSeq All ASVs

**Figure 13.**
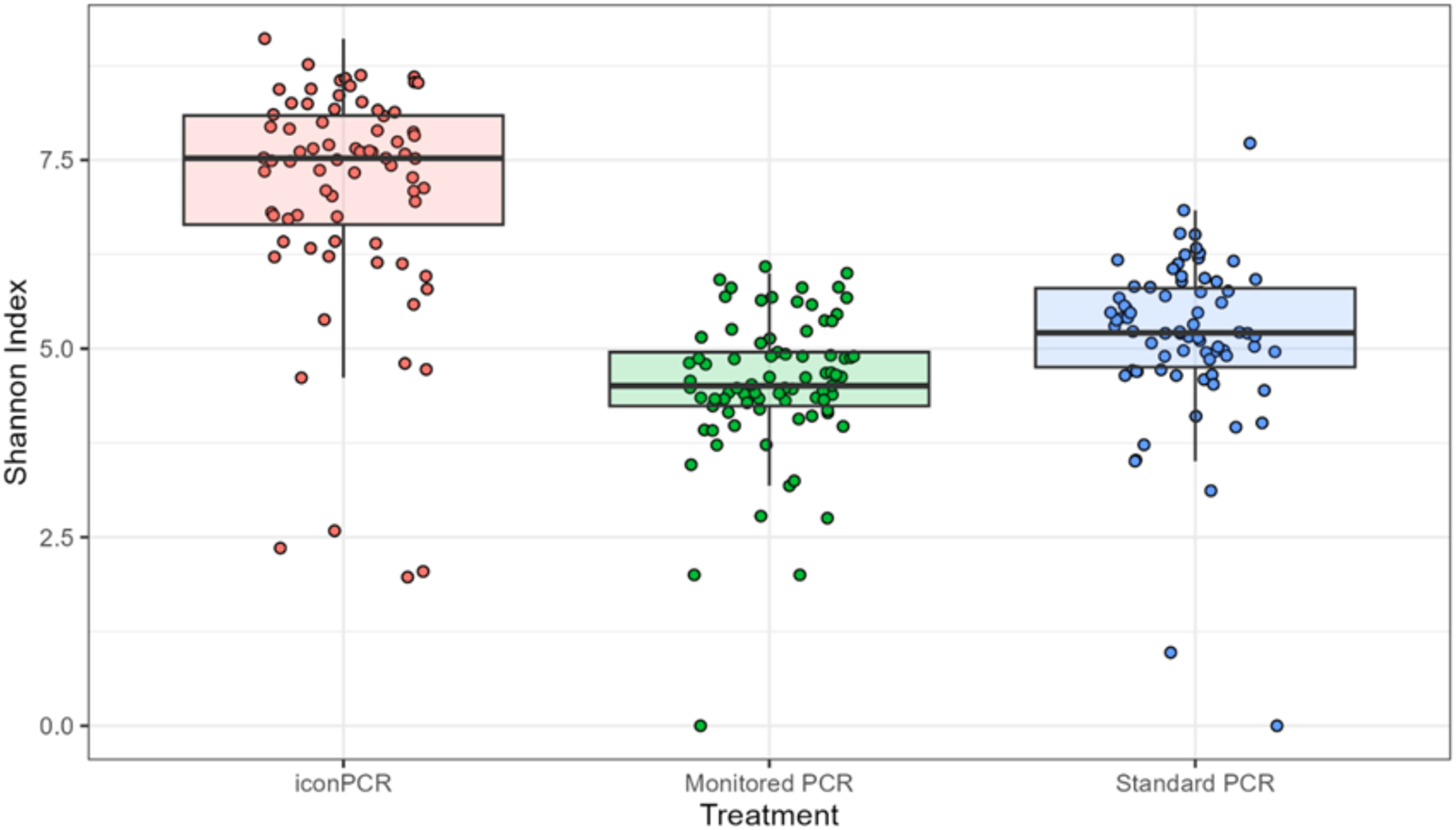
Detected Species Diversity by Treatment - LoopSeq No Singletons

**Figure 14.**
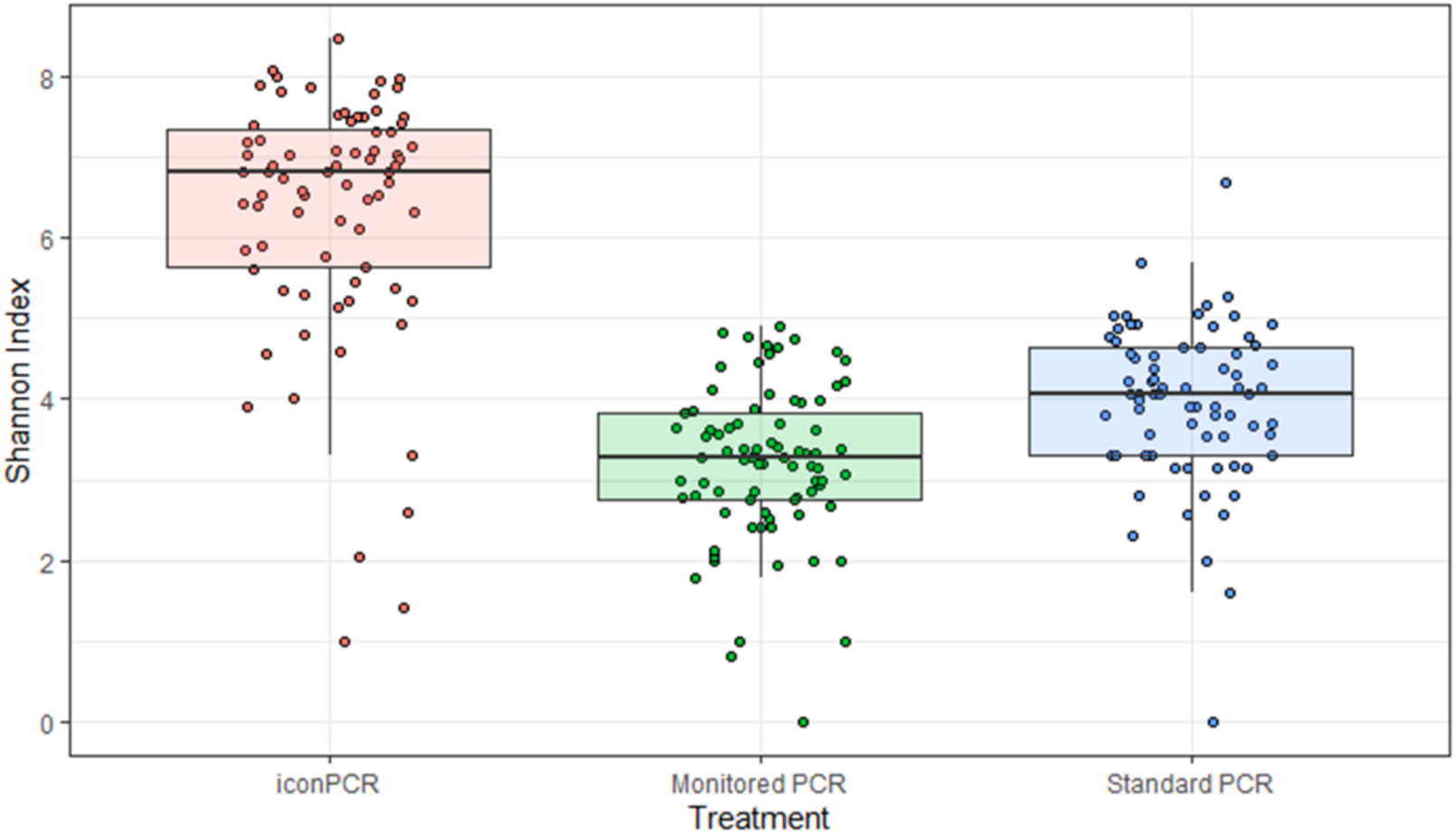
Detected Species Diversity by Treatment - LoopSeq No Doubletons

Species diversity equality was not detected using the Friedman test at 5% significance across all three treatments, nor with the Wilcoxon test at 5% significance between treatment pairs (p<0.0001), regardless of whether singletons or doubletons were excluded. However, Bland-Altman evaluations for method equivalency indicated that Shannon Index values for matched iconPCR and Monitored PCR samples could be considered equivalent (p=0.0429) when all ASVs were included, as well as between Monitored PCR and Standard PCR samples (p<0.0001) when singletons or doubletons were removed with an allowable difference of ± 1.3. Removing singletons and doubletons enhances the discernible detection of sample diversity when the iconPCR in Auto-Normalization mode is employed over other methods by reducing over-amplification.

## Conclusion

The impact of PCR bias on taxa representation has long been known and studied (Suzuki & Giovannoni, 1996). Annealing temperatures and DNA structures are predominant factors contributing to the representation bias (Peng *et al*., 2018). The presence of PCR inhibitors commonly found in environmental samples (Jiang *et al., 2005; Sáenz et al.*, 2019) can contribute to the variability in amplification efficiencies during library preparation. Finally, over-amplification can result in the higher representation of certain taxa compared to others based on their different amplification efficiency, causing species to be obscured or underrepresented. The Auto-Normalization mode of the iconPCR system allows users to limit the number of cycles per sample to only the amount required to achieve a given amplification value corresponding to a theoretical final yield. In doing so, representation bias is minimized, as are sequencing artifacts. While certain amplicons may have a PCR efficiency close to optimal (2-fold factor at each cycle), certain factors (GC content, secondary structures) may reduce the relative representation of other species. After nine cycles, an amplicon with a 1.5-fold replication efficiency will be 10 times less abundant than an amplicon with a 2-fold efficiency. The results presented in this article illustrate the significant reduction in the rate of off-target products (Figures 6 and 7), 1.3 to 4.3-fold reduction in chimeras (Figures 8 and 9), and ∼2-fold increase in the Shannon Diversity Index (Figures 13 - 14) for iconPCR libraries. These findings are consistent for both small (V3 and V4) and large amplicon (V1-V9) generation, the latter of which was confirmed using alternative redundant 16S long-read assays (LoopSeq by Element Biosciences and 16S rRNA by PacBio) and sequencing platforms (AVITI and Sequel IIe). In addition to limiting overamplification, auto-normalization ensures that samples will not be dropped out due to under-amplification when insufficient cycles would not lead to enough material being sequenced properly.

When evaluating the species diversity of a given microbiome, classification using the full V1-V9 region of the 16S rRNA subunit offers a more comprehensive view of the genetic diversity present in a sample, allowing for more accurate taxonomic assignments (Jeong et al., 2021; Buetas et al., 2024). In the LoopSeq assay, synthetic long reads are classified as full-length, regardless of the anticipated length, if the downstream pipeline detects both 5’ and 3’ adapters at the termini of the consensus read (Liu et al., 2021). However, only reads classified as full-length or end-only (in which the 3’ adapter sequence was detected) are subjected to ASV assignment and taxonomic classification. By comparison, all PacBio sequences, regardless of their expected lengths, are considered full-length. For our analyses of V1-V9 molecules, 1276 bps was utilized as the lower cutoff for comparison of all full-length fragments across assays, as shorter molecules were generally not flagged as full-length by the LoopSeq pipeline. This length was in line with that previously used in the analysis of PacBio data (Zhang et al. 2024).

An evaluation of the full-length molecules between 1276-1726 bps in size depicts the critical impact on the observed species diversity within microbial communities when samples are subjected to overamplification (Table 3). Standard PCR amplifications led to a 2-4-fold increased formation of chimeras (Figure 9), potentially artificially inflating the perceived diversity of a sample by introducing spurious sequences (Figures 13 and 14). This phenomenon, previously demonstrated by Polz and Cavanaugh (1998), can obscure the true biological diversity and result in misleading conclusions about community structure. Callahan et al. (2021) emphasize the importance of full-length sequencing in accurately capturing microbial diversity. The integration of shorter sequences into ASV analysis skews the diversity metrics, as these fragments may not represent distinct species or operational taxonomic units (OTUs) but partial or degraded sequences that lack the necessary resolution for accurate identification. As Bunge et al. (2012) highlight, implementing error correction strategies at the initial data generation stage can enhance the reliability of the data by reducing noise. This strategy is employed in the error-correction generation of the consensus synthetic long LoopSeq reads. This practice often leads to identifying more distinct ASVs, thus increasing the Shannon Diversity Index.

Although the more laborious Monitored PCR approach may appear to yield similar diversity metrics, after singleton and doubleton removal, it is apparent that iconPCR provides several benefits that justify its adoption. The work presented here shows that the high-fidelity amplification of full-length molecules minimizes the production of artifacts, such as chimeras, thus ensuring that the reported diversity reflects true biological variation rather than sequencing errors. Furthermore, the capability of iconPCR to produce longer, more accurate reads enhances taxonomic resolution and functional predictions, ultimately leading to better insights into microbial ecology and evolution. iconPCR should, therefore, be viewed as a way to obtain high-quality, reliable data that advances our understanding of complex microbial communities. This study also illustrates the considerable workflow advantages of iconPCR. While it is possible to optimize cycling parameters by monitoring the profiles for each sample, this process is relatively more laborious and time-consuming. It also involves several thermocyclers if samples are to be run simultaneously. By adopting iconPCR, users can improve the quality of their microbiome data while considerably simplifying their sequencing workflow. The workflow improvements also apply to other NGS library prep methods, which are all affected by over- or under-amplification.

