## Supplemental Data for "The use of iconPCR for 16S library preparation improves data quality and workflow"

#### Supplemental Table 1. Short Read Auto-Normalized Cycle Number

| Assigned Index | Sample Name | V3 Plate Well | V3 Cycle Number | V4 Plate Well | V4 Cycle Number |
|---|---|---|---|---|---|
| A1 | Control_Cultivar1_Rep_1_a | A1 | 21 | A1 | 15 |
| A2 | Trt_Cultivar1_Rep_3_a | A2 | 19 | A2 | 13 |
| A3 | Control_Cultivar1_Rep_1_b | A3 | 21 | A3 | 13 |
| A4 | Trt_Cultivar1_Rep_3_b | A4 | 21 | A4 | 14 |
| A5 | Control_Cultivar1_Rep_1_c | A5 | 21 | A5 | 15 |
| A6 | Trt_Cultivar1_Rep_3_c | A6 | 21 | A6 | 15 |
| A7 | Control_Cultivar1_Rep_1_d | A7 | 21 | A7 | 13 |
| A8 | Trt_Cultivar1_Rep_3_d | A8 | 21 | A8 | 13 |
| A9 | Control_T1_1 | A9 | 19 | A9 | 13 |
| A10 | AgriGro_T1_4 | A10 | 19 | A10 | 11 |
| A11 | AgriGro_T2_2 | A11 | 19 | A11 | 11 |
| B1 | Control_Cultivar2_Rep_1_a | B1 | 21 | B1 | 15 |
| B2 | Trt_Cultivar2_Rep_3_a | B2 | 19 | B2 | 12 |
| B3 | Control_Cultivar2_Rep_1_b | B3 | 21 | B3 | 14 |
| B4 | Trt_Cultivar2_Rep_3_b | B4 | 19 | B4 | 13 |
| B5 | Control_Cultivar2_Rep_1_c | B5 | 21 | B5 | 15 |
| B6 | Trt_Cultivar2_Rep_3_c | B6 | 19 | B6 | 14 |
| B7 | Control_Cultivar2_Rep_1_d | B7 | 19 | B7 | 14 |
| B8 | Trt_Cultivar2_Rep_3_d | B8 | 21 | B8 | 14 |
| B9 | Control_T1_2 | B9 | 19 | B9 | 13 |
| B10 | AgriGro_T1_5 | B10 | 19 | B10 | 14 |
| B11 | AgriGro_T2_3 | B11 | 19 | B11 | 13 |
| C1 | TrT_Cultivar1_Rep_1_a | C1 | 21 | C1 | 14 |
| C2 | Control_Cultivar1_Rep_3_a | C2 | 19 | C2 | 13 |
| C3 | TrT_Cultivar1_Rep_1_b | C3 | 19 | C3 | 14 |
| C4 | Control_Cultivar1_Rep_3_b | C4 | 21 | C4 | 14 |
| C5 | TrT_Cultivar1_Rep_1_c | C5 | 21 | C5 | 14 |
| C6 | Control_Cultivar1_Rep_3_c | C6 | 21 | C6 | 14 |

| Assigned Index | Sample Name | V3 Plate Well | V3 Cycle Number | V4 Plate Well | V4 Cycle Number |
|---|---|---|---|---|---|
| C7 | TrT_Cultivar1_Rep_1_d | C7 | 19 | C7 | 13 |
| C8 | Control_Cultivar1_Rep_3_d | C8 | 19 | C8 | 13 |
| C9 | Control_T1_3 | C9 | 19 | C9 | 13 |
| C10 | Control_T2_1 | C10 | 19 | C10 | 12 |
| C11 | AgriGro_T2_4 | C11 | 19 | C11 | 14 |
| D1 | TrT_Cultivar2_Rep_1_a | D1 | 21 | D1 | 14 |
| D2 | Control_Cultivar2_Rep_3_a | D2 | 19 | D2 | 13 |
| D3 | TrT_Cultivar2_Rep_1_b | D3 | 19 | D3 | 13 |
| D4 | Control_Cultivar2_Rep_3_b | D4 | 19 | D4 | 14 |
| D5 | TrT_Cultivar2_Rep_1_c | D5 | 21 | D5 | 14 |
| D6 | Control_Cultivar2_Rep_3_c | D6 | 21 | D6 | 14 |
| D7 | TrT_Cultivar2_Rep_1_d | D7 | 21 | D7 | 14 |
| D8 | Control_Cultivar2_Rep_3_d | D8 | 21 | D8 | 13 |
| D9 | Control_T1_4 | D9 | 19 | D9 | 13 |
| D10 | Control_T2_2 | D10 | 19 | D10 | 13 |
| D11 | AgriGro_T2_5 | D11 | 19 | D11 | 14 |
| E1 | TrT_Cultivar1_Rep_2_a | E1 | 19 | E1 | 13 |
| E2 | Control_Cultivar1_Rep_4_a | E2 | 19 | E2 | 14 |
| E3 | TrT_Cultivar1_Rep_2_b | E3 | 21 | E3 | 14 |
| E4 | Control_Cultivar1_Rep_4_b | E4 | 19 | E4 | 14 |
| E5 | TrT_Cultivar1_Rep_2_c | E5 | 21 | E5 | 16 |
| E6 | Control_Cultivar1_Rep_4_c | E6 | 21 | E6 | 13 |
| E7 | TrT_Cultivar1_Rep_2_d | E7 | 21 | E7 | 14 |
| E8 | Control_Cultivar1_Rep_4_d | E8 | 21 | E8 | 14 |
| E9 | Control_T1_5 | E9 | 19 | E9 | 13 |
| E10 | Control_T2_3 | E10 | 19 | E10 | 14 |
| E11 | Negative Control | E11 | 23 | E11 | 18 |
| F1 | TrT_Cultivar2_Rep_2_a | F1 | 19 | F1 | 13 |
| F2 | Control_Cultivar2_Rep_4_a | F2 | 19 | F2 | 14 |
| F3 | TrT_Cultivar2_Rep_2_b | F3 | 21 | F3 | 13 |
| F4 | Control_Cultivar2_Rep_4_b | F4 | 19 | F4 | 14 |

| Assigned Index | Sample Name | V3 Plate Well | V3 Cycle Number | V4 Plate Well | V4 Cycle Number |
|---|---|---|---|---|---|
| F5 | TrT_Cultivar2_Rep_2_c | F5 | 21 | F5 | 14 |
| F6 | Control_Cultivar2_Rep_4_c | F6 | 21 | F6 | 13 |
| F7 | TrT_Cultivar2_Rep_2_d | F7 | 21 | F7 | 14 |
| F8 | Control_Cultivar2_Rep_4_d | F8 | 21 | F8 | 13 |
| F9 | AgriGro_T1_1 | F9 | 19 | F9 | 12 |
| F10 | Control_T2_4 | F10 | 19 | F10 | 14 |
| F11 | Negative Control | F11 | 23 | F11 | 18 |
| G1 | Control_Cultivar1_Rep_2_a | G1 | 19 | G1 | 12 |
| G2 | Trt_Cultivar1_Rep_4_a | G2 | 19 | G2 | 13 |
| G3 | Control_Cultivar1_Rep_2_b | G3 | 19 | G3 | 13 |
| G4 | Trt_Cultivar1_Rep_4_b | G4 | 21 | G4 | 14 |
| G5 | Control_Cultivar1_Rep_2_c | G5 | 21 | G5 | 13 |
| G6 | Trt_Cultivar1_Rep_4_c | G6 | 21 | G6 | 14 |
| G7 | Control_Cultivar1_Rep_2_d | G7 | 19 | G7 | 14 |
| G8 | Trt_Cultivar1_Rep_4_d | G8 | 23 | G8 | 16 |
| G9 | AgriGro_T1_2 | G9 | 19 | G9 | 13 |
| G10 | Control_T2_5 | G10 | 19 | G10 | 14 |
| H1 | Control_Cultivar2_Rep_2_a | H1 | 19 | H2 | 11 |
| H2 | Trt_Cultivar2_Rep_4_a | H2 | 19 | H3 | 13 |
| H3 | Control_Cultivar2_Rep_2_b | H3 | 19 | H4 | 12 |
| H4 | Trt_Cultivar2_Rep_4_b | H4 | 21 | H5 | 12 |
| H5 | Control_Cultivar2_Rep_2_c | H5 | 19 | H6 | 12 |
| H6 | Trt_Cultivar2_Rep_4_c | H6 | 21 | H7 | 12 |
| H7 | Control_Cultivar2_Rep_2_d | H7 | 19 | H8 | 13 |
| H8 | Trt_Cultivar2_Rep_4_d | H8 | 19 | H9 | 12 |
| H9 | AgriGro_T1_3 | H9 | 19 | H10 | 11 |
| H10 | AgriGro_T2_1 | H10 | 19 | H11 | 13 |

**Supplemental Table 2. Long Read Auto-Normalized Cycle Number**

N/E - Sample Not Evaluated

| Well | Sample Name | LoopSeq | PacBio |
|---|---|---|---|
| A1 | Control_Cultivar1_Rep_1_a | 21 | 19 |
| A2 | Trt_Cultivar1_Rep_3_a | 18 | 19 |
| A3 | Control_Cultivar1_Rep_1_b | 30 | 18 |
| A4 | Trt_Cultivar1_Rep_3_b | 30 | 19 |
| A5 | Control_Cultivar1_Rep_1_c | 30 | 19 |
| A6 | Trt_Cultivar1_Rep_3_c | 30 | 19 |
| A7 | Control_Cultivar1_Rep_1_d | 8 | 18 |
| A8 | Trt_Cultivar1_Rep_3_d | 30 | 18 |
| A9 | Control_T1_1 | 30 | N/E |
| A10 | AgriGro_T1_4 | 30 | N/E |
| A11 | AgriGro_T2_2 | 30 | N/E |
| B1 | Control_Cultivar2_Rep_1_a | 18 | 18 |
| B2 | Trt_Cultivar2_Rep_3_a | 18 | 19 |
| B3 | Control_Cultivar2_Rep_1_b | 17 | 19 |
| B4 | Trt_Cultivar2_Rep_3_b | 17 | 19 |
| B5 | Control_Cultivar2_Rep_1_c | 17 | 20 |
| B6 | Trt_Cultivar2_Rep_3_c | 17 | 19 |
| B7 | Control_Cultivar2_Rep_1_d | 17 | 18 |
| B8 | Trt_Cultivar2_Rep_3_d | 17 | 19 |
| B9 | Control_T1_2 | 18 | N/E |
| B10 | AgriGro_T1_5 | 17 | N/E |
| B11 | AgriGro_T2_3 | 25 | N/E |
| C1 | TrT_Cultivar1_Rep_1_a | 17 | 18 |
| C2 | Control_Cultivar1_Rep_3_a | 18 | 19 |
| C3 | TrT_Cultivar1_Rep_1_b | 20 | 19 |
| C4 | Control_Cultivar1_Rep_3_b | 16 | 20 |
| C5 | TrT_Cultivar1_Rep_1_c | 16 | 20 |
| C6 | Control_Cultivar1_Rep_3_c | 16 | 18 |
| C7 | TrT_Cultivar1_Rep_1_d | 20 | 19 |
| C8 | Control_Cultivar1_Rep_3_d | 17 | 18 |
| C9 | Control_T1_3 | 18 | N/E |
| C10 | Control_T2_1 | 18 | N/E |

| Well | Sample Name | LoopSeq | PacBio |
|---|---|---|---|
| C11 | AgriGro_T2_4 | 18 | N/E |
| D1 | TrT_Cultivar2_Rep_1_a | 18 | 19 |
| D2 | Control_Cultivar2_Rep_3_a | 18 | 18 |
| D3 | TrT_Cultivar2_Rep_1_b | 18 | 17 |
| D4 | Control_Cultivar2_Rep_3_b | 19 | 17 |
| D5 | TrT_Cultivar2_Rep_1_c | 16 | 17 |
| D6 | Control_Cultivar2_Rep_3_c | 16 | 19 |
| D7 | TrT_Cultivar2_Rep_1_d | 17 | 17 |
| D8 | Control_Cultivar2_Rep_3_d | 16 | 17 |
| D9 | Control_T1_4 | 18 | N/E |
| D10 | Control_T2_2 | 18 | N/E |
| D11 | AgriGro_T2_5 | 17 | N/E |
| E1 | TrT_Cultivar1_Rep_2_a | 21 | 19 |
| E2 | Control_Cultivar1_Rep_4_a | 18 | 19 |
| E3 | TrT_Cultivar1_Rep_2_b | 17 | 20 |
| E4 | Control_Cultivar1_Rep_4_b | 18 | 19 |
| E5 | TrT_Cultivar1_Rep_2_c | 7 | 19 |
| E6 | Control_Cultivar1_Rep_4_c | 17 | 18 |
| E7 | TrT_Cultivar1_Rep_2_d | 17 | 18 |
| E8 | Control_Cultivar1_Rep_4_d | 16 | 19 |
| E9 | Control_T1_5 | 17 | N/E |
| E10 | Control_T2_3 | 17 | N/E |
| E11 | Zymo Control | 18 | 14 |
| F1 | TrT_Cultivar2_Rep_2_a | 19 | 19 |
| F2 | Control_Cultivar2_Rep_4_a | 18 | 18 |
| F3 | TrT_Cultivar2_Rep_2_b | 17 | 19 |
| F4 | Control_Cultivar2_Rep_4_b | 18 | 18 |
| F5 | TrT_Cultivar2_Rep_2_c | 17 | 19 |
| F6 | Control_Cultivar2_Rep_4_c | 30 | 18 |
| F7 | TrT_Cultivar2_Rep_2_d | 16 | 17 |
| F8 | Control_Cultivar2_Rep_4_d | 16 | 18 |
| F9 | AgriGro_T1_1 | 20 | N/E |
| F10 | Control_T2_4 | 18 | N/E |
| G1 | Control_Cultivar1_Rep_2_a | 23 | 19 |

| Well | Sample Name | LoopSeq | PacBio |
|---|---|---|---|
| G2 | Trt_Cultivar1_Rep_4_a | 18 | 18 |
| G3 | Control_Cultivar1_Rep_2_b | 21 | 18 |
| G4 | Trt_Cultivar1_Rep_4_b | 16 | 21 |
| G5 | Control_Cultivar1_Rep_2_c | 17 | 18 |
| G6 | Trt_Cultivar1_Rep_4_c | 16 | 18 |
| G7 | Control_Cultivar1_Rep_2_d | 17 | 18 |
| G8 | Trt_Cultivar1_Rep_4_d | 30 | 17 |
| G9 | AgriGro_T1_2 | 18 | N/E |
| G10 | Control_T2_5 | 23 | N/E |
| H2 | Trt_Cultivar2_Rep_4_a | 20 | 18 |
| H3 | Control_Cultivar2_Rep_2_b | 19 | 18 |
| H4 | Trt_Cultivar2_Rep_4_b | 17 | 19 |
| H5 | Control_Cultivar2_Rep_2_c | 17 | 18 |
| H6 | Trt_Cultivar2_Rep_4_c | 17 | 18 |
| H7 | Control_Cultivar2_Rep_2_d | 20 | 17 |
| H8 | Trt_Cultivar2_Rep_4_d | 18 | 19 |
| H9 | AgriGro_T1_3 | 17 | N/E |
| H10 | AgriGro_T2_1 | 18 | N/E |
| H11 | Control_Cultivar2_Rep_2_a | 23 | 19 |

### Supplemental Table 3. V3 Chimera Rate and Chimera Free Reads

Bolded values indicate the Lowest Chimera Rate or Highest Chimera-Free Read percentage.

| Sample Name | iconPCR | | | Standard PCR | | |
|---|---|---|---|---|---|---|
| | Read Number | Chimera Rate | Chimera Free Reads (%) | Read Number | Chimera Rate | Chimera Free Reads (%) |
| AgriGro_T1_1 | 136324 | **0.042216** | **95.80%** | 31146 | 0.076093 | 92.40% |
| AgriGro_T1_2 | 121833 | **0.035877** | **96.40%** | 23342 | 0.059421 | 94.10% |
| AgriGro_T1_3 | 0 | NA | N/C | 8614 | 0.017181 | 98.30% |
| AgriGro_T1_4 | 213690 | **0.040746** | **95.90%** | 45812 | 0.088165 | 91.20% |
| AgriGro_T1_5 | 144051 | **0.041145** | **95.90%** | 27071 | 0.067268 | 93.30% |
| AgriGro_T2_1 | 141892 | **0.043716** | **95.60%** | 35876 | 0.110101 | 89.00% |
| AgriGro_T2_2 | 129193 | **0.030156** | **97.00%** | 58503 | 0.064595 | 93.50% |

| Sample Name | iconPCR | | | Standard PCR | | |
|---|---|---|---|---|---|---|
| | Read Number | Chimera Rate | Chimera Free Reads (%) | Read Number | Chimera Rate | Chimera Free Reads (%) |
| AgriGro_T2_3 | 91888 | **0.023899** | **97.60%** | 26482 | 0.033041 | 96.70% |
| AgriGro_T2_4 | 125550 | **0.038471** | **96.20%** | 79335 | 0.074129 | 92.60% |
| AgriGro_T2_5 | 115720 | **0.042274** | **95.80%** | 49717 | 0.050425 | 95.00% |
| Control_Cultivar1_Rep_1_a | 268915 | **0.041749** | **95.80%** | 54183 | 0.062197 | 93.80% |
| Control_Cultivar1_Rep_1_b | 270628 | **0.059754** | **94.00%** | 36451 | 0.067735 | 93.20% |
| Control_Cultivar1_Rep_1_c | 159212 | **0.031266** | **96.90%** | 22183 | 0.021368 | 97.90% |
| Control_Cultivar1_Rep_1_d | 123291 | **0.02971** | **97.00%** | 45562 | 0.053356 | 94.70% |
| Control_Cultivar1_Rep_2_a | 195464 | **0.025104** | **97.50%** | 62024 | 0.070037 | 93.00% |
| Control_Cultivar1_Rep_2_b | 228157 | **0.030299** | **97.00%** | 60963 | 0.060332 | 94.00% |
| Control_Cultivar1_Rep_2_c | 300342 | 0.045974 | 95.40% | 34723 | **0.041068** | **95.90%** |
| Control_Cultivar1_Rep_2_d | 133025 | **0.028679** | **97.10%** | 25121 | 0.040643 | 95.90% |
| Control_Cultivar1_Rep_3_a | 164683 | **0.019304** | **98.10%** | 87765 | 0.055181 | 94.50% |
| Control_Cultivar1_Rep_3_b | 123903 | **0.022639** | **97.70%** | 37230 | 0.053263 | 94.70% |
| Control_Cultivar1_Rep_3_c | 104400 | **0.025278** | **97.50%** | 28766 | 0.034555 | 96.50% |
| Control_Cultivar1_Rep_3_d | 280278 | 0.039607 | 96.00% | 8137 | **0.013641** | **98.60%** |
| Control_Cultivar1_Rep_4_a | 160334 | **0.014482** | **98.60%** | 49437 | 0.025932 | 97.40% |
| Control_Cultivar1_Rep_4_b | 107686 | **0.013669** | **98.60%** | 61553 | 0.040762 | 95.90% |
| Control_Cultivar1_Rep_4_c | 136358 | **0.023328** | **97.70%** | 38947 | 0.042494 | 95.80% |
| Control_Cultivar1_Rep_4_d | 110054 | 0.019636 | 98.00% | 13279 | **0.015438** | **98.50%** |
| Control_Cultivar2_Rep_1_a | 185876 | **0.03484** | **96.50%** | 35986 | 0.056327 | 94.40% |
| Control_Cultivar2_Rep_1_b | 315847 | **0.05553** | **94.40%** | 61880 | 0.097027 | 90.30% |
| Control_Cultivar2_Rep_1_c | 192509 | **0.040331** | **96.00%** | 39912 | 0.057652 | 94.20% |
| Control_Cultivar2_Rep_1_d | 319304 | **0.043914** | **95.60%** | 29893 | 0.04941 | 95.10% |
| Control_Cultivar2_Rep_2_a | 113585 | **0.01332** | **98.70%** | 51703 | 0.049804 | 95.00% |
| Control_Cultivar2_Rep_2_b | 176467 | **0.025705** | **97.40%** | 37106 | 0.071929 | 92.80% |
| Control_Cultivar2_Rep_2_c | 527533 | 0.052734 | 94.70% | 10275 | **0.018686** | **98.10%** |
| Control_Cultivar2_Rep_2_d | 167968 | **0.021046** | **97.90%** | 39110 | 0.060291 | 94.00% |
| Control_Cultivar2_Rep_3_a | 256970 | **0.028548** | **97.10%** | 85691 | 0.070813 | 92.90% |
| Control_Cultivar2_Rep_3_b | 119238 | **0.017696** | **98.20%** | 82470 | 0.091949 | 90.80% |

| Sample Name | iconPCR | | | Standard PCR | | |
|---|---|---|---|---|---|---|
| | Read Number | Chimera Rate | Chimera Free Reads (%) | Read Number | Chimera Rate | Chimera Free Reads (%) |
| Control_Cultivar2_Rep_3_c | 114222 | **0.023481** | **97.70%** | 29067 | 0.040389 | 96.00% |
| Control_Cultivar2_Rep_3_d | 113429 | **0.023336** | **97.70%** | 24698 | 0.042797 | 95.70% |
| Control_Cultivar2_Rep_4_a | 227159 | **0.023451** | **97.70%** | 52108 | 0.052161 | 94.80% |
| Control_Cultivar2_Rep_4_b | 129056 | **0.01762** | **98.20%** | 43279 | 0.043947 | 95.60% |
| Control_Cultivar2_Rep_4_c | 124451 | **0.024251** | **97.60%** | 17902 | 0.023126 | 97.70% |
| Control_Cultivar2_Rep_4_d | 104540 | 0.01807 | 98.20% | 15897 | **0.012204** | **98.80%** |
| Control_T1_1 | 148173 | **0.031483** | **96.90%** | 28294 | 0.040998 | 95.90% |
| Control_T1_2 | 141154 | **0.035309** | **96.50%** | 26287 | 0.062198 | 93.80% |
| Control_T1_3 | 75936 | **0.016079** | **98.40%** | 16463 | 0.036506 | 96.30% |
| Control_T1_4 | 109673 | **0.023123** | **97.70%** | 28607 | 0.057084 | 94.30% |
| Control_T1_5 | 142774 | **0.026651** | **97.30%** | 45229 | 0.076522 | 92.30% |
| Control_T2_1 | 143922 | **0.034185** | **96.60%** | 23188 | 0.064559 | 93.50% |
| Control_T2_2 | 126548 | **0.037401** | **96.30%** | 62615 | 0.120259 | 88.00% |
| Control_T2_3 | 161925 | **0.043625** | **95.60%** | 22332 | 0.051406 | 94.90% |
| Control_T2_4 | 157005 | **0.033993** | **96.60%** | 46822 | 0.097668 | 90.20% |
| Control_T2_5 | 118696 | **0.045503** | **95.40%** | 38916 | 0.101424 | 89.90% |
| TrT_Cultivar1_Rep_1_a | 138261 | **0.028591** | **97.10%** | 69818 | 0.076284 | 92.40% |
| TrT_Cultivar1_Rep_1_b | 263045 | **0.028968** | **97.10%** | 47478 | 0.051687 | 94.80% |
| TrT_Cultivar1_Rep_1_c | 266870 | **0.038768** | **96.10%** | 35364 | 0.047591 | 95.20% |
| TrT_Cultivar1_Rep_1_d | 138008 | **0.023999** | **97.60%** | 44953 | 0.061153 | 93.90% |
| TrT_Cultivar1_Rep_2_a | 188678 | **0.017336** | **98.30%** | 48081 | 0.042428 | 95.80% |
| TrT_Cultivar1_Rep_2_b | 206217 | **0.031297** | **96.90%** | 39660 | 0.051563 | 94.80% |
| TrT_Cultivar1_Rep_2_c | 201279 | **0.035061** | **96.50%** | 54077 | 0.058676 | 94.10% |
| TrT_Cultivar1_Rep_2_d | 123520 | **0.031776** | **96.80%** | 49259 | 0.061958 | 93.80% |
| Trt_Cultivar1_Rep_3_a | 249085 | **0.03066** | **96.90%** | 77627 | 0.071354 | 92.90% |
| Trt_Cultivar1_Rep_3_b | 216489 | **0.035822** | **96.40%** | 83849 | 0.079428 | 92.10% |
| Trt_Cultivar1_Rep_3_c | 265282 | **0.036961** | **96.30%** | 85146 | 0.06685 | 93.30% |
| Trt_Cultivar1_Rep_3_d | 237080 | **0.035992** | **96.40%** | 27050 | 0.037745 | 96.20% |
| Trt_Cultivar1_Rep_4_a | 105956 | **0.012373** | **98.80%** | 49318 | 0.030597 | 96.90% |

| Sample Name | iconPCR | | | Standard PCR | | |
|---|---|---|---|---|---|---|
| | Read Number | Chimera Rate | Chimera Free Reads (%) | Read Number | Chimera Rate | Chimera Free Reads (%) |
| Trt_Cultivar1_Rep_4_b | 240777 | **0.036548** | **96.30%** | 32704 | 0.037824 | 96.20% |
| Trt_Cultivar1_Rep_4_c | 80859 | **0.015014** | **98.50%** | 43280 | 0.039718 | 96.00% |
| Trt_Cultivar1_Rep_4_d | 185197 | **0.068252** | **93.20%** | 71413 | 0.087925 | 91.20% |
| TrT_Cultivar2_Rep_1_a | 217189 | **0.031894** | **96.80%** | 43066 | 0.056913 | 94.30% |
| TrT_Cultivar2_Rep_1_b | 240398 | **0.030574** | **96.90%** | 48807 | 0.066138 | 93.40% |
| TrT_Cultivar2_Rep_1_c | 277254 | **0.067891** | **93.20%** | 78331 | 0.220602 | 77.90% |
| TrT_Cultivar2_Rep_1_d | 123960 | **0.030937** | **96.90%** | 25275 | 0.045579 | 95.40% |
| TrT_Cultivar2_Rep_2_a | 172232 | **0.01916** | **98.10%** | 31976 | 0.047067 | 95.30% |
| TrT_Cultivar2_Rep_2_b | 184364 | **0.036905** | **96.30%** | 29854 | 0.053594 | 94.60% |
| TrT_Cultivar2_Rep_2_c | 161047 | **0.029141** | **97.10%** | 54656 | 0.070788 | 92.90% |
| TrT_Cultivar2_Rep_2_d | 121951 | **0.027249** | **97.30%** | 29486 | 0.042596 | 95.70% |
| Trt_Cultivar2_Rep_3_a | 190092 | **0.020643** | **97.90%** | 38976 | 0.046105 | 95.40% |
| Trt_Cultivar2_Rep_3_b | 183960 | **0.035481** | **96.50%** | 85402 | 0.108124 | 89.20% |
| Trt_Cultivar2_Rep_3_c | 0 | NA | N/C | 26846 | 0.056917 | 94.30% |
| Trt_Cultivar2_Rep_3_d | 319835 | 0.042416 | 95.80% | **5855** | **0.017933** | **98.20%** |
| Trt_Cultivar2_Rep_4_a | 228114 | **0.023449** | **97.70%** | 49445 | 0.04233 | 95.80% |
| Trt_Cultivar2_Rep_4_b | 0 | NA | N/C | 43718 | 0.037079 | 96.30% |
| Trt_Cultivar2_Rep_4_c | 126918 | 0.025962 | 97.40% | 16181 | **0.024968** | **97.50%** |
| Trt_Cultivar2_Rep_4_d | 101902 | 0.023385 | 97.70% | 11824 | **0.007104** | **99.30%** |

### Supplemental Table 4. V4 Chimera Rate and Chimera Free Reads

Bolded values indicate the Lowest Chimera Rate or Highest Chimera-Free Read percentage.

| Sample Name | iconPCR | | | Standard PCR | | |
|---|---|---|---|---|---|---|
| | Read Number | Chimera Rate | Chimera Free Reads (%) | Read Number | Chimera Rate | Chimera Free Reads (%) |
| AgriGro_T1_1 | 1720 | **0.113953** | **88.60%** | 45333 | 0.331855 | 66.80% |
| AgriGro_T1_2 | 1383 | **0.236443** | **76.40%** | 63781 | 0.412129 | 58.80% |
| AgriGro_T1_3 | 5142 | **0.271879** | **72.80%** | 85149 | 0.550106 | 45.00% |
| AgriGro_T1_4 | 264 | **0** | **100.00%** | 38421 | 0.364879 | 63.50% |

| Sample Name | iconPCR | | | Standard PCR | | |
|---|---|---|---|---|---|---|
| | Read Number | Chimera Rate | Chimera Free Reads (%) | Read Number | Chimera Rate | Chimera Free Reads (%) |
| AgriGro_T1_5 | 8820 | **0.081406** | **91.90%** | 63528 | 0.489485 | 51.10% |
| AgriGro_T2_1 | 462 | **0.002165** | **99.80%** | 34437 | 0.19752 | 80.20% |
| AgriGro_T2_2 | 742 | **0.033693** | **96.60%** | 8524 | 0.363679 | 63.60% |
| AgriGro_T2_3 | 2117 | **0.011337** | **98.90%** | 53471 | 0.430402 | 57.00% |
| AgriGro_T2_4 | 1028 | **0.015564** | **98.40%** | 23072 | 0.351205 | 64.90% |
| AgriGro_T2_5 | 1382 | **0.013025** | **98.70%** | 18367 | 0.363042 | 63.70% |
| Control_Cultivar1_Rep_1_a | 1987 | **0.054857** | **94.50%** | 57800 | 0.439637 | 56.00% |
| Control_Cultivar1_Rep_1_b | 4372 | **0.351784** | **64.80%** | 136249 | 0.610449 | 39.00% |
| Control_Cultivar1_Rep_1_c | 893 | **0** | **100.00%** | 39296 | 0.512011 | 48.80% |
| Control_Cultivar1_Rep_1_d | 1382 | **0.002171** | **99.80%** | 74206 | 0.276864 | 72.30% |
| Control_Cultivar1_Rep_2_a | 558 | **0.09319** | **90.70%** | 50300 | 0.400159 | 60.00% |
| Control_Cultivar1_Rep_2_b | 1144 | **0.227273** | **77.30%** | 81695 | 0.41262 | 58.70% |
| Control_Cultivar1_Rep_2_c | 1567 | **0.199745** | **80.00%** | 75326 | 0.426081 | 57.40% |
| Control_Cultivar1_Rep_2_d | 1410 | **0.109929** | **89.00%** | 36016 | 0.477982 | 52.20% |
| Control_Cultivar1_Rep_3_a | 3459 | **0.011275** | **98.90%** | 57345 | 0.330752 | 66.90% |
| Control_Cultivar1_Rep_3_b | 663 | **0** | **100.00%** | 42464 | 0.437265 | 56.30% |
| Control_Cultivar1_Rep_3_c | 1456 | **0.03228** | **96.80%** | 14194 | 0.298225 | 70.20% |
| Control_Cultivar1_Rep_3_d | 999 | **0** | **100.00%** | 27476 | 0.315039 | 68.50% |
| Control_Cultivar1_Rep_4_a | 1148 | **0.143728** | **85.60%** | 70225 | 0.452873 | 54.70% |
| Control_Cultivar1_Rep_4_b | 1681 | **0.267103** | **73.30%** | 58134 | 0.545619 | 45.40% |
| Control_Cultivar1_Rep_4_c | 1338 | **0.022422** | **97.80%** | 6314 | 0.384701 | 61.50% |
| Control_Cultivar1_Rep_4_d | 838 | **0.23747** | **76.30%** | 8496 | 0.387123 | 61.30% |
| Control_Cultivar2_Rep_1_a | 1974 | **0.291793** | **70.80%** | 86012 | 0.487897 | 51.20% |
| Control_Cultivar2_Rep_1_b | 1660 | **0.086145** | **91.40%** | 58609 | 0.360559 | 63.90% |
| Control_Cultivar2_Rep_1_c | 951 | **0.001052** | **99.90%** | 89755 | 0.407164 | 59.30% |
| Control_Cultivar2_Rep_1_d | 3936 | **0.454268** | **54.60%** | 18390 | 0.714628 | 28.50% |
| Control_Cultivar2_Rep_2_a | 1576 | **0** | **100.00%** | 86530 | 0.426384 | 57.40% |
| Control_Cultivar2_Rep_2_b | 1720 | **0** | **100.00%** | 85026 | 0.369452 | 63.10% |
| Control_Cultivar2_Rep_2_c | 671 | **0.086438** | **91.40%** | 145066 | 0.276047 | 72.40% |

| Sample Name | iconPCR | | | Standard PCR | | |
|---|---|---|---|---|---|---|
| | Read Number | Chimera Rate | Chimera Free Reads (%) | Read Number | Chimera Rate | Chimera Free Reads (%) |
| Control_Cultivar2_Rep_2_d | 2345 | **0.074627** | **92.50%** | 37488 | 0.44833 | 55.20% |
| Control_Cultivar2_Rep_3_a | 7985 | **0.404634** | **59.50%** | 147493 | 0.58737 | 41.30% |
| Control_Cultivar2_Rep_3_b | 3385 | **0.114328** | **88.60%** | 29623 | 0.413463 | 58.70% |
| Control_Cultivar2_Rep_3_c | 1403 | **0.021383** | **97.90%** | 57534 | 0.247749 | 75.20% |
| Control_Cultivar2_Rep_3_d | 818 | **0.135697** | **86.40%** | 46994 | 0.280078 | 72.00% |
| Control_Cultivar2_Rep_4_a | 3357 | **0.251713** | **74.80%** | 111485 | 0.581271 | 41.90% |
| Control_Cultivar2_Rep_4_b | 2819 | **0.260731** | **73.90%** | 53671 | 0.434667 | 56.50% |
| Control_Cultivar2_Rep_4_c | 1646 | **0.108141** | **89.20%** | 8667 | 0.467174 | 53.30% |
| Control_Cultivar2_Rep_4_d | 739 | **0.104195** | **89.60%** | 34806 | 0.32888 | 67.10% |
| Control_T1_1 | 1598 | **0.003755** | **99.60%** | 93180 | 0.258146 | 74.20% |
| Control_T1_2 | 1765 | **0.002833** | **99.70%** | 73445 | 0.297679 | 70.20% |
| Control_T1_3 | 1302 | **0.147465** | **85.30%** | 2299 | 0.254023 | 74.60% |
| Control_T1_4 | 1272 | **0.112421** | **88.80%** | 10395 | 0.408754 | 59.10% |
| Control_T1_5 | 1192 | **0** | **100.00%** | 65955 | 0.340399 | 66.00% |
| Control_T2_1 | 1072 | **0.069963** | **93.00%** | 48211 | 0.281741 | 71.80% |
| Control_T2_2 | 1283 | **0.039751** | **96.00%** | 44636 | 0.480509 | 51.90% |
| Control_T2_3 | 3490 | **0.280802** | **71.90%** | 28446 | 0.47961 | 52.00% |
| Control_T2_4 | 1688 | **0.033768** | **96.60%** | 40631 | 0.440476 | 56.00% |
| Control_T2_5 | 4139 | **0.043247** | **95.70%** | 6305 | 0.227914 | 77.20% |
| TrT_Cultivar1_Rep_1_a | 924 | **0** | **100.00%** | 51453 | 0.306785 | 69.30% |
| TrT_Cultivar1_Rep_1_b | 1398 | **0.11588** | **88.40%** | 23078 | 0.35796 | 64.20% |
| TrT_Cultivar1_Rep_1_c | 802 | **0** | **100.00%** | 72936 | 0.404862 | 59.50% |
| TrT_Cultivar1_Rep_1_d | 1909 | **0.190676** | **80.90%** | 35274 | 0.426603 | 57.30% |
| TrT_Cultivar1_Rep_2_a | 1703 | **0.058133** | **94.20%** | 38603 | 0.285703 | 71.40% |
| TrT_Cultivar1_Rep_2_b | 2607 | **0.13809** | **86.20%** | 49462 | 0.439004 | 56.10% |
| TrT_Cultivar1_Rep_2_c | 2333 | **0.190742** | **80.90%** | 50675 | 0.403039 | 59.70% |
| TrT_Cultivar1_Rep_2_d | 1044 | **0.070881** | **92.90%** | 35931 | 0.409173 | 59.10% |
| Trt_Cultivar1_Rep_3_a | 211 | **0** | **100.00%** | 108034 | 0.384768 | 61.50% |
| Trt_Cultivar1_Rep_3_b | 913 | **0.066813** | **93.30%** | 30102 | 0.359245 | 64.10% |

| Sample Name | iconPCR | | | Standard PCR | | |
|---|---|---|---|---|---|---|
| | Read Number | Chimera Rate | Chimera Free Reads (%) | Read Number | Chimera Rate | Chimera Free Reads (%) |
| Trt_Cultivar1_Rep_3_c | 1177 | **0.002549** | **99.70%** | 165587 | 0.339912 | 66.00% |
| Trt_Cultivar1_Rep_3_d | 1732 | **0.034065** | **96.60%** | 91058 | 0.394035 | 60.60% |
| Trt_Cultivar1_Rep_4_a | 3050 | **0.11541** | **88.50%** | 115798 | 0.511175 | 48.90% |
| Trt_Cultivar1_Rep_4_b | 1230 | **0.000813** | **99.90%** | 37213 | 0.369199 | 63.10% |
| Trt_Cultivar1_Rep_4_c | 1267 | **0.149171** | **85.10%** | 47711 | 0.359707 | 64.00% |
| Trt_Cultivar1_Rep_4_d | 972 | **0.280864** | **71.90%** | 39969 | 0.432936 | 56.70% |
| TrT_Cultivar2_Rep_1_a | 1087 | **0.093836** | **90.60%** | 125452 | 0.401835 | 59.80% |
| TrT_Cultivar2_Rep_1_b | 1466 | **0.002729** | **99.70%** | 5336 | 0.380435 | 62.00% |
| TrT_Cultivar2_Rep_1_c | 515 | **0.01165** | **98.80%** | 47688 | 0.385653 | 61.40% |
| TrT_Cultivar2_Rep_1_d | 1890 | **0.060847** | **93.90%** | 60988 | 0.368007 | 63.20% |
| TrT_Cultivar2_Rep_2_a | 1949 | **0.278091** | **72.20%** | 52835 | 0.436358 | 56.40% |
| TrT_Cultivar2_Rep_2_b | 1807 | **0.016602** | **98.30%** | 50410 | 0.247768 | 75.20% |
| TrT_Cultivar2_Rep_2_c | 883 | **0** | **100.00%** | 24755 | 0.370713 | 62.90% |
| TrT_Cultivar2_Rep_2_d | 1188 | **0.083333** | **91.70%** | 17080 | 0.47096 | 52.90% |
| Trt_Cultivar2_Rep_3_a | 2052 | **0.079922** | **92.00%** | 30651 | 0.386382 | 61.40% |
| Trt_Cultivar2_Rep_3_b | 1092 | **0** | **100.00%** | 106196 | 0.409488 | 59.10% |
| Trt_Cultivar2_Rep_3_c | 1436 | **0.000696** | **99.90%** | 48459 | 0.402526 | 59.70% |
| Trt_Cultivar2_Rep_3_d | 914 | **0** | **100.00%** | 33785 | 0.326891 | 67.30% |
| Trt_Cultivar2_Rep_4_a | 2348 | **0** | **100.00%** | 47773 | 0.292948 | 70.70% |
| Trt_Cultivar2_Rep_4_b | 443 | **0** | **100.00%** | 38508 | 0.349694 | 65.00% |
| Trt_Cultivar2_Rep_4_c | 1184 | **0.032939** | **96.70%** | 75443 | 0.504275 | 49.60% |
| Trt_Cultivar2_Rep_4_d | 671 | **0** | **100.00%** | 33621 | 0.557955 | 44.20% |

### Supplemental Table 5. LoopSeq Long Read Chimeric Rate and Chimera Free Reads

N/C - Not Calculated

| Sample | Full-Length Molecules | | | Chimera Rate | | | Chimera Free Reads (%) | | |
|---|---|---|---|---|---|---|---|---|---|
| | Monitored PCR | iconPCR | Standard PCR | Monitored PCR | iconPCR | Standard PCR | Monitored PCR | iconPCR | Standard PCR |
| AgriGro_T1_1 | 7657 | 5394 | 6983 | 0 | 0 | 0.003030303 | 100.0% | 100.0% | 99.7% |

| Sample | Full-Length Molecules | | | Chimera Rate | | | Chimera Free Reads (%) | | |
|---|---|---|---|---|---|---|---|---|---|
| | Monitored PCR | iconPCR | Standard PCR | Monitored PCR | iconPCR | Standard PCR | Monitored PCR | iconPCR | Standard PCR |
| AgriGro_T1_2 | 6146 | 5824 | 6048 | 0 | 0 | 0.013329848 | 100.0% | 100.0% | 98.7% |
| AgriGro_T1_3 | 5054 | 5165 | 5060 | 0.026315789 | 0 | 0.009911141 | 97.4% | 100.0% | 99.0% |
| AgriGro_T1_4 | 4380 | 973 | 1598 | 0 | 0.029354207 | 0 | 100.0% | 97.1% | 100.0% |
| AgriGro_T1_5 | 3736 | 5225 | 2762 | 0 | 0 | 0 | 100.0% | 100.0% | 100.0% |
| AgriGro_T2_1 | 6081 | 4967 | 1557 | 0 | 0 | 0 | 100.0% | 100.0% | 100.0% |
| AgriGro_T2_2 | 4304 | 4 | 4 | 0 | 0 | 0 | 100.0% | 100.0% | 100.0% |
| AgriGro_T2_3 | 5512 | 1513 | 2569 | 0.017391304 | 0 | 0 | 98.3% | 100.0% | 100.0% |
| AgriGro_T2_4 | 3925 | 1699 | 1370 | 0 | 0 | N/C | 100.0% | 100.0% | N/C |
| AgriGro_T2_5 | 3517 | 3473 | 2179 | 0 | 0 | 0 | 100.0% | 100.0% | 100.0% |
| Control_Cultivar1_Rep_1_a | 4341 | 371 | 835 | 0 | 0 | 0 | 100.0% | 100.0% | 100.0% |
| Control_Cultivar1_Rep_1_b | 4098 | 0 | 3826 | 0 | N/C | 0.000828958 | 100.0% | N/C | 99.9% |
| Control_Cultivar1_Rep_1_c | 3021 | 8 | 928 | 0 | 0 | 0 | 100.0% | 100.0% | 100.0% |
| Control_Cultivar1_Rep_1_d | 3811 | 0 | 444 | 0 | N/C | 0 | 100.0% | N/C | 100.0% |
| Control_Cultivar1_Rep_2_a | 4395 | 131 | 1013 | 0 | 0 | 0 | 100.0% | 100.0% | 100.0% |
| Control_Cultivar1_Rep_2_b | 6951 | 4478 | 3965 | 0 | 0 | 0 | 100.0% | 100.0% | 100.0% |
| Control_Cultivar1_Rep_2_c | 4401 | 4281 | 2246 | 0 | 0 | 0.022113022 | 100.0% | 100.0% | 97.8% |
| Control_Cultivar1_Rep_2_d | 4987 | 2905 | 2940 | 0 | 0 | 0.044117647 | 100.0% | 100.0% | 95.6% |
| Control_Cultivar1_Rep_3_a | 2693 | 5500 | 2400 | 0 | 0 | 0 | 100.0% | 100.0% | 100.0% |
| Control_Cultivar1_Rep_3_b | 4441 | 4979 | 2437 | 0 | 0 | 0 | 100.0% | 100.0% | 100.0% |
| Control_Cultivar1_Rep_3_c | 5547 | 6154 | 3760 | 0 | 0 | 0 | 100.0% | 100.0% | 100.0% |
| Control_Cultivar1_Rep_3_d | 6205 | 8175 | 4417 | 0 | 0 | 0 | 100.0% | 100.0% | 100.0% |
| Control_Cultivar1_Rep_4_a | 4658 | 4815 | 2403 | 0 | 0 | 0 | 100.0% | 100.0% | 100.0% |

| Sample | Full-Length Molecules | | | Chimera Rate | | | Chimera Free Reads (%) | | |
|---|---|---|---|---|---|---|---|---|---|
| | Monitored PCR | iconPCR | Standard PCR | Monitored PCR | iconPCR | Standard PCR | Monitored PCR | iconPCR | Standard PCR |
| Control_Cultivar1_Rep_4_b | 5716 | 7049 | 3802 | 0 | 0 | 0.006166495 | 100.0% | 100.0% | 99.4% |
| Control_Cultivar1_Rep_4_c | 6450 | 4560 | 4569 | 0 | 0 | 0 | 100.0% | 100.0% | 100.0% |
| Control_Cultivar1_Rep_4_d | 5119 | 7086 | 5 | 0 | 0 | 0 | 100.0% | 100.0% | 100.0% |
| Control_Cultivar2_Rep_1_a | 5878 | 4018 | 2541 | 0 | 0 | 0.005836576 | 100.0% | 100.0% | 99.4% |
| Control_Cultivar2_Rep_1_b | 4478 | 6335 | 2684 | 0 | 0 | 0 | 100.0% | 100.0% | 100.0% |
| Control_Cultivar2_Rep_1_c | 3667 | 3693 | 2622 | 0 | 0 | 0 | 100.0% | 100.0% | 100.0% |
| Control_Cultivar2_Rep_1_d | 3121 | 4157 | 1471 | 0 | 0 | 0 | 100.0% | 100.0% | 100.0% |
| Control_Cultivar2_Rep_2_a | 3635 | 1274 | 5129 | 0 | 0 | 0.005214368 | 100.0% | 100.0% | 99.5% |
| Control_Cultivar2_Rep_2_b | 5572 | 6263 | 4310 | 0 | 0 | 0 | 100.0% | 100.0% | 100.0% |
| Control_Cultivar2_Rep_2_c | 4989 | 3260 | 1939 | 0 | 0 | 0 | 100.0% | 100.0% | 100.0% |
| Control_Cultivar2_Rep_2_d | 7180 | 2832 | 4040 | 0 | 0 | 0 | 100.0% | 100.0% | 100.0% |
| Control_Cultivar2_Rep_3_a | 4026 | 5722 | 3006 | 0 | 0 | 0 | 100.0% | 100.0% | 100.0% |
| Control_Cultivar2_Rep_3_b | 5624 | 6051 | 3923 | 0 | 0 | 0 | 100.0% | 100.0% | 100.0% |
| Control_Cultivar2_Rep_3_c | 5744 | 6149 | 2794 | 0 | 0 | 0 | 100.0% | 100.0% | 100.0% |
| Control_Cultivar2_Rep_3_d | 5406 | 7858 | 3935 | 0 | 0 | 0.005415162 | 100.0% | 100.0% | 99.5% |
| Control_Cultivar2_Rep_4_a | 4860 | 6522 | 1883 | 0 | 0 | 0 | 100.0% | 100.0% | 100.0% |
| Control_Cultivar2_Rep_4_b | 5921 | 7737 | 4855 | 0 | 0 | 0.012116317 | 100.0% | 100.0% | 98.8% |
| Control_Cultivar2_Rep_4_c | 185 | 1994 | 8418 | 0 | 0 | 0.006410256 | 100.0% | 100.0% | 99.4% |
| Control_Cultivar2_Rep_4_d | 8848 | 9083 | 12674 | 0 | 0 | 0.007820928 | 100.0% | 100.0% | 99.2% |
| Control_T1_1 | 4612 | 2879 | 7 | 0 | 0.019060284 | 0 | 100.0% | 98.1% | 100.0% |

| Sample | Full-Length Molecules | | | Chimera Rate | | | Chimera Free Reads (%) | | |
|---|---|---|---|---|---|---|---|---|---|
| | Monitored PCR | iconPCR | Standard PCR | Monitored PCR | iconPCR | Standard PCR | Monitored PCR | iconPCR | Standard PCR |
| Control_T1_2 | 4769 | 5345 | 4965 | 0 | 0 | 0 | 100.0% | 100.0% | 100.0% |
| Control_T1_3 | 7341 | 8870 | 4260 | 0 | 0 | 0.006797583 | 100.0% | 100.0% | 99.3% |
| Control_T1_4 | 6100 | 6697 | 4551 | 0 | 0 | 0.020011435 | 100.0% | 100.0% | 98.0% |
| Control_T1_5 | 6707 | 12692 | 6794 | 0 | 0 | 0.017323275 | 100.0% | 100.0% | 98.3% |
| Control_T2_1 | 6420 | 5804 | 3327 | 0 | 0 | 0 | 100.0% | 100.0% | 100.0% |
| Control_T2_2 | 6437 | 9622 | 4537 | 0 | 0.003659652 | 0 | 100.0% | 99.6% | 100.0% |
| Control_T2_3 | 4816 | 6068 | 5471 | 0 | 0 | 0.01556568 | 100.0% | 100.0% | 98.4% |
| Control_T2_4 | 5702 | 5053 | 5345 | 0.046875 | 0 | 0.022134387 | 95.3% | 100.0% | 97.8% |
| Control_T2_5 | 7859 | 5402 | 5231 | 0.032407407 | 0 | 0.019437695 | 96.8% | 100.0% | 98.1% |
| TrT_Cultivar1_ Rep_1_a | 4914 | 4938 | 2402 | 0 | 0 | 0 | 100.0% | 100.0% | 100.0% |
| TrT_Cultivar1_ Rep_1_b | 5866 | 5356 | 2973 | 0 | 0 | 0.029279279 | 100.0% | 100.0% | 97.1% |
| TrT_Cultivar1_ Rep_1_c | 4368 | 4880 | 2986 | 0 | 0 | 0.033248082 | 100.0% | 100.0% | 96.7% |
| TrT_Cultivar1_ Rep_1_d | 6323 | 5588 | 5125 | 0 | 0 | 0.003757633 | 100.0% | 100.0% | 99.6% |
| TrT_Cultivar1_ Rep_2_a | 6158 | 2473 | 2324 | 0 | 0 | 0 | 100.0% | 100.0% | 100.0% |
| TrT_Cultivar1_ Rep_2_b | 3500 | 4417 | 3122 | 0 | 0 | 0 | 100.0% | 100.0% | 100.0% |
| TrT_Cultivar1_ Rep_2_c | 5129 | 0 | 4885 | 0 | N/C | 0 | 100.0% | N/C | 100.0% |
| TrT_Cultivar1_ Rep_2_d | 6378 | 7482 | 4785 | 0 | 0 | 0 | 100.0% | 100.0% | 100.0% |
| Trt_Cultivar1_ Rep_3_a | 4009 | 3627 | 3693 | 0 | 0 | 0 | 100.0% | 100.0% | 100.0% |
| Trt_Cultivar1_ Rep_3_b | 4142 | 4 | 1768 | 0 | 0 | 0.006451613 | 100.0% | 100.0% | 99.4% |
| Trt_Cultivar1_ Rep_3_c | 4887 | 0 | 1267 | 0 | N/C | 0 | 100.0% | N/C | 100.0% |
| Trt_Cultivar1_ Rep_3_d | 3433 | 6 | 81 | 0 | 0 | 0.002267574 | 100.0% | 100.0% | 99.8% |
| Trt_Cultivar1_ Rep_4_a | 6993 | 8983 | 2851 | 0 | 0 | 0 | 100.0% | 100.0% | 100.0% |

| Sample | Full-Length Molecules | | | Chimera Rate | | | Chimera Free Reads (%) | | |
|---|---|---|---|---|---|---|---|---|---|
| | Monitored PCR | iconPCR | Standard PCR | Monitored PCR | iconPCR | Standard PCR | Monitored PCR | iconPCR | Standard PCR |
| Trt_Cultivar1_ Rep_4_b | 4496 | 4119 | 2003 | 0 | 0 | 0 | 100.0% | 100.0% | 100.0% |
| Trt_Cultivar1_ Rep_4_c | 5661 | 3498 | 1740 | 0 | 0 | 0 | 100.0% | 100.0% | 100.0% |
| Trt_Cultivar1_ Rep_4_d | 8589 | 327 | 50 | 0 | 0 | 0 | 100.0% | 100.0% | 100.0% |
| TrT_Cultivar2_ Rep_1_a | 5739 | 4101 | 1446 | 0 | 0 | 0.028239203 | 100.0% | 100.0% | 97.2% |
| TrT_Cultivar2_ Rep_1_b | 7033 | 8693 | 4360 | 0 | 0 | 0.024303497 | 100.0% | 100.0% | 97.6% |
| TrT_Cultivar2_ Rep_1_c | 4771 | 7811 | 2928 | 0.008695652 | 0.054111033 | 0 | 99.1% | 94.6% | 100.0% |
| TrT_Cultivar2_ Rep_1_d | 6496 | 7088 | 4365 | 0 | 0 | 0 | 100.0% | 100.0% | 100.0% |
| TrT_Cultivar2_ Rep_2_a | 589 | 4594 | 2364 | 0 | 0 | 0 | 100.0% | 100.0% | 100.0% |
| TrT_Cultivar2_ Rep_2_b | 5807 | 6362 | 4673 | 0 | 0 | 0 | 100.0% | 100.0% | 100.0% |
| TrT_Cultivar2_ Rep_2_c | 6033 | 7175 | 4994 | 0 | 0 | 0 | 100.0% | 100.0% | 100.0% |
| TrT_Cultivar2_ Rep_2_d | 1 | 7556 | 3617 | 0 | 0 | 0 | 100.0% | 100.0% | 100.0% |
| Trt_Cultivar2_ Rep_3_a | 4869 | 8839 | 3843 | 0 | 0 | 0.007797271 | 100.0% | 100.0% | 99.2% |
| Trt_Cultivar2_ Rep_3_b | 4855 | 5527 | 2378 | 0 | 0 | 0.021978022 | 100.0% | 100.0% | 97.8% |
| Trt_Cultivar2_ Rep_3_c | 3972 | 3587 | 1542 | 0 | 0 | 0 | 100.0% | 100.0% | 100.0% |
| Trt_Cultivar2_ Rep_3_d | 4653 | 4518 | 2400 | 0 | 0 | 0 | 100.0% | 100.0% | 100.0% |
| Trt_Cultivar2_ Rep_4_a | 5132 | 1726 | 992 | 0 | 0 | 0 | 100.0% | 100.0% | 100.0% |
| Trt_Cultivar2_ Rep_4_b | 3487 | 3095 | 2732 | 0 | 0 | 0.007712082 | 100.0% | 100.0% | 99.2% |
| Trt_Cultivar2_ Rep_4_c | 56 | 3142 | 2486 | 0 | 0 | 0 | 100.0% | 100.0% | 100.0% |
| Trt_Cultivar2_ Rep_4_d | 5659 | 3374 | 3201 | 0 | 0 | 0 | 100.0% | 100.0% | 100.0% |
| Zymo | 2748 | 3098 | 1072 | 0.213465953 | 0.141089109 | 0 | 78.7% | 85.9% | 100.0% |

### Supplemental Table 6. PacBio Chimera Free Reads

| Sample | icon PCR | | Standard PCR | | % Chimera Free Reads | |
|---|---|---|---|---|---|---|
| | Trimmed Seq | Chimera Free Seq | Trimmed Seq | Chimera Free Seq | iconPCR | Standard PCR |
| Control_Cultivar1_Rep_1_c | 50273 | 35616 | 11800 | 682 | **70.8%** | 5.8% |
| Control_Cultivar1_Rep_1_d | 56173 | 40704 | 17128 | 2483 | **72.5%** | 14.5% |
| Control_Cultivar1_Rep_2_c | 39938 | 26826 | 10552 | 722 | **67.2%** | 6.8% |
| Control_Cultivar1_Rep_2_d | 43555 | 29761 | 16566 | 2643 | **68.3%** | 16.0% |
| Control_Cultivar1_Rep_3_c | 57987 | 40574 | 11766 | 924 | **70.0%** | 7.9% |
| Control_Cultivar1_Rep_3_d | 34948 | 22979 | 13756 | 2151 | **65.8%** | 15.6% |
| Control_Cultivar1_Rep_4_c | 57016 | 43538 | 31718 | 19354 | **76.4%** | 61.0% |
| Control_Cultivar1_Rep_4_d | 62388 | 43116 | 30101 | 18114 | **69.1%** | 60.2% |
| Control_Cultivar2_Rep_1_c | 32162 | 22045 | 13033 | 1945 | **68.5%** | 14.9% |
| Control_Cultivar2_Rep_1_d | 35569 | 23249 | 19323 | 4310 | **65.4%** | 22.3% |
| Control_Cultivar2_Rep_2_c | 25786 | 15530 | 11493 | 1157 | **60.2%** | 10.1% |
| Control_Cultivar2_Rep_2_d | 25352 | 17028 | 21103 | 6175 | **67.2%** | 29.3% |
| Control_Cultivar2_Rep_3_c | 37072 | 23299 | 10998 | 1017 | **62.8%** | 9.2% |
| Control_Cultivar2_Rep_3_d | 31615 | 19522 | 10372 | 982 | **61.7%** | 9.5% |
| Control_Cultivar2_Rep_4_c | 41606 | 27171 | 11005 | 504 | **65.3%** | 4.6% |
| Control_Cultivar2_Rep_4_d | 46494 | 30476 | 10947 | 606 | **65.5%** | 5.5% |
| TrT_Cultivar1_ Rep_1_c | 78963 | 60802 | 9930 | 310 | **77.0%** | 3.1% |
| TrT_Cultivar1_ Rep_1_d | 41970 | 29756 | 22995 | 6585 | **70.9%** | 28.6% |
| TrT_Cultivar1_ Rep_2_c | 48564 | 38637 | 34494 | 25255 | **79.6%** | 73.2% |
| TrT_Cultivar1_ Rep_2_d | 37271 | 24092 | 59904 | 43962 | 64.6% | **73.4%** |
| Trt_Cultivar1_ Rep_3_c | 28514 | 18340 | 10455 | 516 | **64.3%** | 4.9% |
| Trt_Cultivar1_ Rep_3_d | 29244 | 18568 | 13140 | 1751 | **63.5%** | 13.3% |
| Trt_Cultivar1_ Rep_4_c | 33338 | 20327 | 11458 | 650 | **61.0%** | 5.7% |
| Trt_Cultivar1_ Rep_4_d | 72688 | 62339 | 22629 | 14951 | **85.8%** | 66.1% |
| TrT_Cultivar2_ Rep_1_c | 28388 | 22012 | 15230 | 8002 | **77.5%** | 52.5% |
| TrT_Cultivar2_ Rep_1_d | 32210 | 21088 | 14891 | 2510 | **65.5%** | 16.9% |
| TrT_Cultivar2_ Rep_2_c | 40773 | 26993 | 9855 | 423 | **66.2%** | 4.3% |
| TrT_Cultivar2_ Rep_2_d | 53978 | 37854 | 13696 | 1200 | **70.1%** | 8.8% |
| Trt_Cultivar2_ Rep_3_c | 26892 | 16489 | 15203 | 2483 | **61.3%** | 16.3% |

| Sample | icon PCR | | Standard PCR | | % Chimera Free Reads | |
|---|---|---|---|---|---|---|
| | Trimmed Seq | Chimera Free Seq | Trimmed Seq | Chimera Free Seq | iconPCR | Standard PCR |
| Trt_Cultivar2_ Rep_3_d | 27389 | 17080 | 11487 | 1944 | **62.4%** | 16.9% |
| Trt_Cultivar2_ Rep_4_c | 36893 | 22823 | 11352 | 1108 | **61.9%** | 9.8% |
| Trt_Cultivar2_ Rep_4_d | 71922 | 52159 | 14444 | 2134 | **72.5%** | 14.8% |
| Zymo | 40778 | 40695 | 17480 | 10789 | **99.8%** | 61.7% |

### Supplemental Table 7. Shannon Diversity Values - All ASVs

N/C - Value Not Calculated
N/E - Sample Not Evaluated
**Highest Diversity** when at least two values are available

| Sample | LoopSeq | | | PacBio | |
|---|---|---|---|---|---|
| | Monitored PCR | iconPCR | Standard PCR | iconPCR | Standard PCR |
| AgriGro_T1_1 | **12.89638825** | 11.69035 | 10.57972701 | N/E | N/E |
| AgriGro_T1_2 | **12.57830962** | 11.83878 | 9.904051462 | N/E | N/E |
| AgriGro_T1_3 | **12.30083559** | 11.72435 | 9.981200134 | N/E | N/E |
| AgriGro_T1_4 | **12.09397543** | 7.490762 | 7.129283017 | N/E | N/E |
| AgriGro_T1_5 | **11.85830931** | 11.72217 | 9.81856825 | N/E | N/E |
| AgriGro_T2_1 | **12.56445817** | 11.71467 | 8.549866487 | N/E | N/E |
| AgriGro_T2_2 | **12.06507069** | 0 | 1 | N/E | N/E |
| AgriGro_T2_3 | **12.4191521** | 9.542925 | 9.075240459 | N/E | N/E |
| AgriGro_T2_4 | **11.92707442** | 10.03098 | 8.516579922 | N/E | N/E |
| AgriGro_T2_5 | **11.77209304** | 11.00496 | 9.133223443 | N/E | N/E |
| Control_Cultivar1_Rep_1_a | **12.0793174** | 7.761549 | 4.523561956 | N/E | N/E |
| Control_Cultivar1_Rep_1_b | **11.9987521** | N/C | 1.584962501 | N/E | N/E |
| Control_Cultivar1_Rep_1_c | 11.56014844 | N/C | N/C | 9.928 | N/C |
| Control_Cultivar1_Rep_1_d | **11.89063648** | N/C | 0 | 10.086 | N/C |
| Control_Cultivar1_Rep_2_a | **12.09965544** | 6.442943 | 8.977803305 | N/E | N/E |
| Control_Cultivar1_Rep_2_b | **12.75818397** | 11.56442 | 10.50310025 | N/E | N/E |
| Control_Cultivar1_Rep_2_c | **12.10026303** | 11.43774 | 9.825380719 | 9.871 | N/C |
| Control_Cultivar1_Rep_2_d | **12.27964332** | 10.88787 | 10.21294438 | 9.933 | N/C |
| Control_Cultivar1_Rep_3_a | 11.38570793 | **11.76492** | 10.04249269 | N/E | N/E |

| Sample | LoopSeq | | | PacBio | |
|---|---|---|---|---|---|
| | **Monitored PCR** | **iconPCR** | **Standard PCR** | **iconPCR** | **Standard PCR** |
| Control_Cultivar1_Rep_3_b | **12.11396676** | 11.66308 | 9.857302931 | N/E | N/E |
| Control_Cultivar1_Rep_3_c | **12.43519259** | 11.88911 | 10.55190655 | 10.207 | N/C |
| Control_Cultivar1_Rep_3_d | **12.59047018** | 12.19219 | 10.62673148 | 9.808 | N/C |
| Control_Cultivar1_Rep_4_a | **12.18205997** | 11.57771 | 9.868623632 | N/E | N/E |
| Control_Cultivar1_Rep_4_b | **12.47750908** | 12.08262 | 10.52614643 | N/E | N/E |
| Control_Cultivar1_Rep_4_c | **12.65112844** | 11.43851 | 10.81892837 | **7.326** | 7.181 |
| Control_Cultivar1_Rep_4_d | **12.31969278** | 12.05637 | 0 | **10.302** | 6.744 |
| Control_Cultivar2_Rep_1_a | **12.51711002** | 11.24636 | 9.848448427 | N/E | N/E |
| Control_Cultivar2_Rep_1_b | **12.12143366** | 11.8435 | 10.12337815 | N/E | N/E |
| Control_Cultivar2_Rep_1_c | **11.83540373** | 11.09311 | 9.946751951 | 9.41 | N/C |
| Control_Cultivar2_Rep_1_d | **11.59071984** | 11.20739 | 9.288879605 | **9.618** | 7.769 |
| Control_Cultivar2_Rep_2_a | **11.82182224** | 9.714883 | 9.351394491 | N/E | N/E |
| Control_Cultivar2_Rep_2_b | **12.44025469** | 11.89754 | 9.962315986 | N/E | N/E |
| Control_Cultivar2_Rep_2_c | **12.28333231** | 11.08207 | 9.721103402 | 9.361 | N/C |
| Control_Cultivar2_Rep_2_d | **12.78826345** | 10.98399 | 10.32832602 | **9.083** | 7.44 |
| Control_Cultivar2_Rep_3_a | **11.96997624** | 11.77235 | 10.00137066 | N/E | N/E |
| Control_Cultivar2_Rep_3_b | **12.44781928** | 11.92261 | 10.39880745 | N/E | N/E |
| Control_Cultivar2_Rep_3_c | **12.48470633** | 11.91912 | 10.15988543 | 9.754 | N/C |
| Control_Cultivar2_Rep_3_d | **12.39465677** | 12.17073 | 10.6163491 | 9.666 | N/C |
| Control_Cultivar2_Rep_4_a | **12.24427146** | 12.06549 | 9.908872119 | N/E | N/E |
| Control_Cultivar2_Rep_4_b | **12.52664122** | 12.33763 | 10.88750963 | N/E | N/E |
| Control_Cultivar2_Rep_4_c | N/C | 10.23129 | **11.5580434** | 9.916 | N/C |
| Control_Cultivar2_Rep_4_d | **13.10062261** | 12.29306 | 11.92306424 | 10.068 | N/C |
| Control_T1_1 | **12.16201273** | 7.757093 | 1 | N/E | N/E |
| Control_T1_2 | **12.21527732** | 11.63649 | 10.65041347 | N/E | N/E |
| Control_T1_3 | **12.83470811** | 12.39992 | 10.44711064 | N/E | N/E |
| Control_T1_4 | **12.57053535** | 12.0939 | 10.55962454 | N/E | N/E |
| Control_T1_5 | 12.70597181 | **12.91825** | 10.53202744 | N/E | N/E |
| Control_T2_1 | **12.64500715** | 11.89187 | 9.961430354 | N/E | N/E |

| Sample | LoopSeq | | | PacBio | |
|---|---|---|---|---|---|
| | Monitored PCR | iconPCR | Standard PCR | iconPCR | Standard PCR |
| Control_T2_2 | **12.63997308** | 12.47328 | 10.18925459 | N/E | N/E |
| Control_T2_3 | **12.2258311** | 11.85371 | 10.381889 | N/E | N/E |
| Control_T2_4 | **12.4743139** | 11.68689 | 10.43969226 | N/E | N/E |
| Control_T2_5 | **12.92875657** | 11.68312 | 10.05598437 | N/E | N/E |
| TrT_Cultivar1_ Rep_1_a | **12.26105414** | 11.6278 | 9.84501265 | N/E | N/E |
| TrT_Cultivar1_ Rep_1_b | **12.51462319** | 11.74175 | 10.23098415 | N/E | N/E |
| TrT_Cultivar1_ Rep_1_c | **12.09000989** | 11.5823 | 10.37468349 | 10.302 | N/C |
| TrT_Cultivar1_ Rep_1_d | **12.61192098** | 11.64818 | 10.31322377 | **9.868** | 8.333 |
| TrT_Cultivar1_ Rep_2_a | **12.58255931** | 10.59896 | 9.49137012 | N/E | N/E |
| TrT_Cultivar1_ Rep_2_b | **11.77313921** | 11.40879 | 9.807945873 | N/E | N/E |
| TrT_Cultivar1_ Rep_2_c | **12.32012092** | N/C | 10.77448918 | 6.604 | **7.349** |
| TrT_Cultivar1_ Rep_2_d | **12.63657498** | 12.26247 | 10.81513362 | **9.772** | 7.13 |
| Trt_Cultivar1_ Rep_3_a | **11.96703119** | 11.35555 | 10.11407722 | N/E | N/E |
| Trt_Cultivar1_ Rep_3_b | **12.00639077** | N/C | 0.918295834 | N/E | N/E |
| Trt_Cultivar1_ Rep_3_c | 12.24813125 | N/C | N/C | 9.558 | N/C |
| Trt_Cultivar1_ Rep_3_d | **11.73920844** | N/C | 0 | 9.518 | N/C |
| Trt_Cultivar1_ Rep_4_a | **12.76611599** | 12.43977 | 9.999178255 | N/E | N/E |
| Trt_Cultivar1_ Rep_4_b | **12.13114454** | 11.37467 | 10.30864789 | N/E | N/E |
| Trt_Cultivar1_ Rep_4_c | **12.46198168** | 11.1957 | 9.312769891 | 9.702 | N/C |
| Trt_Cultivar1_ Rep_4_d | N/C | N/C | N/C | **4.705** | 2.037 |
| TrT_Cultivar2_ Rep_1_a | **12.48109319** | 11.32044 | 9.15205651 | N/E | N/E |
| TrT_Cultivar2_ Rep_1_b | **12.76932741** | 12.30335 | 10.5317402 | N/E | N/E |
| TrT_Cultivar2_ Rep_1_c | **12.19275038** | 11.75871 | 9.850126479 | **7.129** | 4.412 |
| TrT_Cultivar2_ Rep_1_d | **12.65944542** | 12.05085 | 10.82968907 | 9.524 | N/C |
| TrT_Cultivar2_ Rep_2_a | 9.202123824 | **11.40399** | 9.529389621 | N/E | N/E |
| TrT_Cultivar2_ Rep_2_b | **12.49724791** | 11.91039 | 10.60087942 | N/E | N/E |
| TrT_Cultivar2_ Rep_2_c | **12.554763** | 12.13634 | 10.36715902 | 9.812 | N/C |
| TrT_Cultivar2_ Rep_2_d | 0 | **12.03541** | 10.42487094 | 10.152 | N/C |
| Trt_Cultivar2_ Rep_3_a | 12.24458132 | **12.3936** | 10.14278123 | N/E | N/E |

| Sample | LoopSeq | | | PacBio | |
|---|---|---|---|---|---|
| | Monitored PCR | iconPCR | Standard PCR | iconPCR | Standard PCR |
| Trt_Cultivar2_ Rep_3_b | **12.23778603** | 11.69322 | 9.678994693 | N/E | N/E |
| Trt_Cultivar2_ Rep_3_c | **11.95243871** | 11.0667 | 9.110736381 | 9.248 | N/C |
| Trt_Cultivar2_ Rep_3_d | **12.17555958** | 11.45305 | 9.858649905 | 9.444 | N/C |
| Trt_Cultivar2_ Rep_4_a | **12.32282009** | 10.16482 | 7.059575997 | N/E | N/E |
| Trt_Cultivar2_ Rep_4_b | **11.76777065** | 11.06808 | 10.08598207 | N/E | N/E |
| Trt_Cultivar2_ Rep_4_c | 5.807354922 | **11.02032** | 9.634098248 | 9.775 | N/C |
| Trt_Cultivar2_ Rep_4_d | **12.46301725** | 11.12287 | 10.35378183 | 10.361 | N/C |
| Zymo | **10.89261167** | 9.569822 | 8.390397161 | 4.305 | **4.389** |

### Supplemental Table 8. Shannon Diversity Values No Singletons - LoopSeq

N/C - Value Not Calculated
**Highest Diversity** when at least two values are available

| Sample | No Singletons | | | No Doubletons | | |
|---|---|---|---|---|---|---|
| | Monitored PCR | iconPCR | Standard PCR | Monitored PCR | iconPCR | Standard PCR |
| AgriGro_T1_1 | 4.441092818 | **6.214392631** | 6.124460599 | 3.343354621 | **4.939822782** | 4.939530081 |
| AgriGro_T1_2 | 4.623217841 | **6.767723955** | 5.378019208 | 2.996641329 | **5.464661719** | 3.894224895 |
| AgriGro_T1_3 | 3.721928095 | **6.749338047** | 4.642434649 | 0 | **5.305239787** | 3.307188576 |
| AgriGro_T1_4 | **4.680757756** | 1.970950594 | N/C | **2.75** | 1 | N/C |
| AgriGro_T1_5 | 4.239097918 | **6.764283766** | 5.144923109 | 2.780639062 | **5.61326969** | 3.985228136 |
| AgriGro_T2_1 | 4.406821129 | **6.141865845** | 3.72491954 | 2.858458593 | **4.79365747** | 2.55605421 |
| AgriGro_T2_2 | 4.066784213 | N/C | N/C | 2.556656707 | N/C | N/C |
| AgriGro_T2_3 | 4.18386772 | 4.803609964 | **5.473852295** | 1.950063756 | 3.311278124 | **4.43486528** |
| AgriGro_T2_4 | 4.652569011 | **5.585838627** | 4.10390991 | 2.748940047 | **4.554229297** | 3.308270835 |
| AgriGro_T2_5 | 4.321721231 | **6.395632283** | 4.696321466 | 2.983458593 | **5.379678655** | 3.155020907 |
| Control_Cultivar1_Rep_1_a | **4.390319531** | 2.355388542 | N/C | **3.272905595** | 2.046439345 | N/C |
| Control_Cultivar1_Rep_1_b | 3.921928095 | N/C | N/C | 2.807354922 | N/C | N/C |
| Control_Cultivar1_Rep_1_c | 3.969815782 | N/C | N/C | 3.169925001 | N/C | N/C |
| Control_Cultivar1_Rep_1_d | 4.863291943 | N/C | N/C | 4.226474118 | N/C | N/C |
| Control_Cultivar1_Rep_2_a | **3.182005815** | 2.584962501 | 3.115455557 | 2.405639062 | **2.584962501** | 2.570190637 |
| Control_Cultivar1_Rep_2_b | 5.807009421 | **6.223537647** | 5.476380893 | 4.456564762 | **5.208500041** | 4.155153137 |

| Sample | No Singletons | | | No Doubletons | | |
|---|---|---|---|---|---|---|
| | Monitored PCR | iconPCR | Standard PCR | Monitored PCR | iconPCR | Standard PCR |
| Control_Cultivar1_Rep_2_c | 2.753434386 | **7.331821168** | 4.711019011 | 0.811278124 | **6.832292228** | 2.794238586 |
| Control_Cultivar1_Rep_2_d | 5.45552554 | **7.088758033** | 5.202877111 | 4.396291529 | **6.53014516** | 3.308508892 |
| Control_Cultivar1_Rep_3_a | 2.779950001 | **7.869262224** | 5.401007844 | 2 | **7.310530135** | 4.066779655 |
| Control_Cultivar1_Rep_3_b | 3.913977073 | **8.001202664** | 5.023633651 | 2.521640636 | **7.310822185** | 4.208464386 |
| Control_Cultivar1_Rep_3_c | 4.481727679 | **8.254506056** | 5.888024176 | 2.846439345 | **7.576993662** | 4.236912542 |
| Control_Cultivar1_Rep_3_d | 5.256168855 | **8.524421009** | 6.52740353 | 3.848884617 | **7.862902148** | 5.27077821 |
| Control_Cultivar1_Rep_4_a | 4.616874606 | **7.609661508** | 4.966430732 | 3.277613437 | **7.023671147** | 3.681127543 |
| Control_Cultivar1_Rep_4_b | 3.979097891 | **8.160607623** | 5.823135244 | 2.405639062 | **7.433150223** | 4.570132187 |
| Control_Cultivar1_Rep_4_c | 4.879664005 | **7.580259951** | 6.267767619 | 4.110577243 | **6.907944029** | 4.549531857 |
| Control_Cultivar1_Rep_4_d | 4.315824334 | **8.482996058** | N/C | 2.947702779 | **7.865030589** | N/C |
| Control_Cultivar2_Rep_1_a | 4.395998871 | **5.789689067** | 5.216144745 | 3.405639062 | **5.14571137** | 3.885491389 |
| Control_Cultivar2_Rep_1_b | 4.33743546 | **8.134509788** | 5.698299815 | 3.26633264 | **7.519098635** | 5.029486724 |
| Control_Cultivar2_Rep_1_c | 4.155045547 | **7.483453212** | 5.75020466 | 2.788450457 | **6.896360207** | 4.72220962 |
| Control_Cultivar2_Rep_1_d | 4.675743324 | **7.608452542** | 4.64301016 | 3.618633047 | **6.832265144** | 3.685815709 |
| Control_Cultivar2_Rep_2_a | 3.246439345 | **4.72438566** | 4.526303971 | 1 | **4.006238929** | 3.549787382 |
| Control_Cultivar2_Rep_2_b | 4.351409766 | **7.890440229** | 5.3191319 | 3.572469459 | **7.193705467** | 4.292591231 |
| Control_Cultivar2_Rep_2_c | 4.106603137 | **7.095998558** | 3.95869497 | 3.169925001 | **6.480615155** | 2.799007754 |
| Control_Cultivar2_Rep_2_d | 5.91382331 | **6.421713544** | 4.446650042 | 4.779814998 | **5.770983142** | 3.528240039 |
| Control_Cultivar2_Rep_3_a | 4.898153435 | **7.701109343** | 5.106788228 | 3.970175521 | **7.049535436** | 4.218024566 |
| Control_Cultivar2_Rep_3_b | 4.464661719 | **7.652741483** | 5.81422195 | 3.355834261 | **6.989514074** | 4.772630107 |
| Control_Cultivar2_Rep_3_c | 4.871928095 | **8.583176234** | 5.202901085 | 3.700439718 | **7.937963344** | 3.896240625 |
| Control_Cultivar2_Rep_3_d | 4.900229543 | **8.767624368** | 5.959671134 | 3.337175341 | **8.07046772** | 4.628994789 |
| Control_Cultivar2_Rep_4_a | 4.506890596 | **8.085373235** | 4.976143834 | 3.25 | **7.510045707** | 3.79326969 |
| Control_Cultivar2_Rep_4_b | 5.132944045 | **8.624678914** | 5.610051539 | 3.641604168 | **7.974545147** | 4.149141423 |
| Control_Cultivar2_Rep_4_c | N/C | 6.417976674 | **6.834152944** | N/C | **5.90819393** | 5.701630565 |
| Control_Cultivar2_Rep_4_d | 5.689765206 | **9.108913601** | 7.723291172 | 4.064392248 | **8.466801478** | 6.687716264 |
| Control_T1_1 | **5.372142236** | 2.046439345 | N/C | **3.997952699** | 1.405639062 | N/C |
| Control_T1_2 | 4.869747127 | **7.366267468** | 6.511769145 | 3.616348566 | **6.330694487** | 5.029534161 |
| Control_T1_3 | 5.674747626 | **8.269714774** | 5.406796869 | 4.577567157 | **7.085164136** | 3.685815709 |
| Control_T1_4 | 5.36530336 | **7.651870529** | 4.957660708 | 3.683542362 | **6.542991181** | 4.054128884 |

| Sample | No Singletons | | | No Doubletons | | |
|---|---|---|---|---|---|---|
| | Monitored PCR | iconPCR | Standard PCR | Monitored PCR | iconPCR | Standard PCR |
| Control_T1_5 | 5.151017645 | **8.442410994** | 5.933929559 | 3.456564762 | **7.452616788** | 4.892369275 |
| Control_T2_1 | 4.572431251 | **6.71798049** | 4.85683676 | 3.530493057 | **5.839158151** | 4.074164743 |
| Control_T2_2 | 5.813390201 | **8.35769502** | 6.058197418 | 4.574875776 | **7.404345033** | 5.156989871 |
| Control_T2_3 | 5.621440429 | **7.620455652** | 5.163441436 | 4.628108095 | **6.578610731** | 3.572735872 |
| Control_T2_4 | 4.476409766 | **7.268586557** | 5.568766555 | 3.188721876 | **6.227430503** | 3.794238586 |
| Control_T2_5 | 6.087428599 | **6.330406609** | 5.478439004 | 4.826151804 | **5.222445383** | 4.546970944 |
| TrT_Cultivar1_Rep_1_a | 3.725480557 | **7.427918764** | 5.157962016 | 2.584962501 | **6.753013343** | 3.307188576 |
| TrT_Cultivar1_Rep_1_b | 4.415458134 | **7.128800465** | 5.541353021 | 3.188721876 | **6.318084288** | 4.513894935 |
| TrT_Cultivar1_Rep_1_c | 4.310443058 | **8.103829725** | 5.20107044 | 2 | **7.501363655** | 4.380430488 |
| TrT_Cultivar1_Rep_1_d | 5.68093663 | **7.350670552** | 6.160970288 | 3.83418372 | **6.530220933** | 4.931434234 |
| TrT_Cultivar1_Rep_2_a | 4.197159723 | **5.383898919** | 4.651407829 | 3.141635408 | **4.601838196** | 3.16310135 |
| TrT_Cultivar1_Rep_2_b | 2 | **7.824372862** | 4.905003028 | 2 | **7.212949308** | 3.155913095 |
| TrT_Cultivar1_ Rep_2_c | 4.953416562 | N/C | **6.244533269** | 3.990601563 | N/C | **4.765604068** |
| TrT_Cultivar1_Rep_2_d | 4.810672622 | **8.601898383** | 5.671067153 | 3.892407119 | **4.601838196** | 4.373097974 |
| Trt_Cultivar1_Rep_3_a | 4.521640636 | **6.126111815** | 3.506890596 | 3 | **7.212949308** | 1.584962501 |
| Trt_Cultivar1_Rep_3_b | **4.792982038** | N/C | 0 | 3.391892645 | N/C | N/C |
| Trt_Cultivar1_Rep_3_c | 5.640920652 | N/C | N/C | 4.664578374 | N/C | N/C |
| Trt_Cultivar1_ Rep_3_d | 4.898153435 | N/C | N/C | 3.323231429 | N/C | N/C |
| Trt_Cultivar1_Rep_4_a | 4.624751985 | **7.940842553** | 5.073526956 | 3.370950594 | **7.040268558** | 4.073174882 |
| Trt_Cultivar1_Rep_4_b | 4.415061012 | **7.504130399** | 4.015311532 | 1.79248125 | **6.965056872** | 2.307188576 |
| Trt_Cultivar1_Rep_4_c | 4.925523369 | **7.740412702** | 4.719852359 | 3.642149882 | **7.077780931** | 2.799007754 |
| Trt_Cultivar1_Rep_4_d | N/C | N/C | N/C | N/C | N/C | N/C |
| TrT_Cultivar2_Rep_1_a | 4.47745927 | **6.805022179** | 3.523231429 | 3.057476076 | **6.119596191** | 1.985228136 |
| TrT_Cultivar2_Rep_1_b | 5.808615365 | **8.55499538** | 6.175058998 | 4.740601563 | **7.791415128** | 4.940358415 |
| TrT_Cultivar2_Rep_1_c | 4.910330117 | **7.521375967** | 5.91760343 | 2.669534154 | **6.671460385** | 4.669407256 |
| TrT_Cultivar2_ Rep_1_d | 5.582644981 | **8.175211383** | 6.333138659 | 4.490919498 | **7.549215153** | 5.072655564 |
| TrT_Cultivar2_Rep_2_a | 0 | **7.52594953** | 4.949692895 | N/C | **6.825953982** | 3.894224895 |
| TrT_Cultivar2_Rep_2_b | 4.148955904 | **8.245923789** | 6.199794984 | 2.046439345 | **7.510001202** | 4.875137964 |
| TrT_Cultivar2_Rep_2_c | 4.331976695 | **8.437044313** | 5.29856294 | 2.594906618 | **7.809359839** | 3.570190637 |
| TrT_Cultivar2_Rep_2_d | N/C | **8.531765727** | 5.893776528 | N/C | **7.889565313** | 5.029486724 |

| Sample | No Singletons | | | No Doubletons | | |
|---|---|---|---|---|---|---|
| | Monitored PCR | iconPCR | Standard PCR | Monitored PCR | iconPCR | Standard PCR |
| Trt_Cultivar2_Rep_3_a | 4.331976695 | **7.490740161** | 5.761734795 | 2.968918564 | **6.695933392** | 4.633206219 |
| Trt_Cultivar2_Rep_3_b | 5.232728193 | **7.913513329** | 5.02747316 | 4.164735179 | **7.145959615** | 4.155020907 |
| Trt_Cultivar2_ Rep_3_c | 4.282484261 | **7.601082744** | 4.697325709 | 2.846439345 | **7.035063942** | 3.307188576 |
| Trt_Cultivar2_Rep_3_d | 5.07191335 | **7.604198891** | 5.22229394 | 3.370950594 | **6.909780994** | 4.066370599 |
| Trt_Cultivar2_Rep_4_a | 3.46132014 | **4.613858096** | 0.970950594 | 2.405639062 | **3.896291529** | 0 |
| Trt_Cultivar2_ Rep_4_b | 2 | **7.02329091** | 4.590820101 | 1 | **6.417206173** | 3.155913095 |
| Trt_Cultivar2_ Rep_4_c | N/C | **6.950376412** | 4.974057528 | N/C | **6.393811655** | 3.146539862 |
| Trt_Cultivar2_ Rep_4_d | 4.349199939 | **7.529336747** | 5.227240697 | 2.128085279 | **6.815983239** | 4.155265821 |
| Zymo | **6.000351207** | 5.961008561 | 4.897948922 | 4.910371081 | **5.337112845** | 3.793014649 |
